# Flexibility in the social structure of male chimpanzees (*Pan troglodytes schweinfurthii*) in the Budongo Forest, Uganda

**DOI:** 10.1101/2021.12.11.472209

**Authors:** Gal Badihi, Kelsey Bodden, Klaus Zuberbühler, Liran Samuni, Catherine Hobaiter

## Abstract

Individuals of social species face a trade-off between the competitive costs and social benefits of group living. Species show a range of social strategies to deal with this trade-off, for example atomistic fission-fusion dynamics in which temporary social groups of varying size and membership form and re-form; or molecular fission-fusion dynamics which contain stable sets of multilevel nested subgroups. Chimpanzees are considered an archetypical atomistic fission-fusion species, using dynamic changes in day-to-day association to moderate the costs of within-group competition. It has been argued that humans’ highly flexible social organisation allows us to live in extremely large groups. Using four years of association data from two neighbouring communities of East African chimpanzees (*Pan troglodytes schweinfurthii*), we describe new levels of flexibility in chimpanzee social organisation and confirm the presence of subgrouping in a second, large community of chimpanzees. We show that males from the larger Waibira community (N males 24-31) exhibited additional levels of semi-stable subgrouping, while males from the smaller Sonso community (N males 10-13) did not. Subgroup membership showed stability across some years, but flexibility across others. Our data support the hypothesis that chimpanzees can incorporate strategies other than fission-fusion to overcome costs of social living, and that their social organisation may be closer to that of modern humans than previously described.

**SIGNIFICANCE STATEMENT:** Social living offers benefits and costs; groups can more easily locate and defend resources, but experience increased individual competition. Many species, or individuals, flexibly adjust their social organization when faced with different competitive pressures. It is argued that humans are unique among primates in combining multigroup social organisation with fission-fusion dynamics flexibly within and across groups, and that doing so allows us to live in extremely large groups. Using four-years of association data we show new levels of flexibility in chimpanzee social organization. Males from a typically-sized community employed atomistic fission-fusion dynamics, but males in an unusually large community incorporated additional semi-stable levels of subgrouping. Our data support the hypothesis that chimpanzee males combine social organization strategies, and that doing so may allow them, like humans, to mitigate individual costs at larger community sizes.

## INTRODUCTION

Social animals derive extensive benefits from group living, including maintenance of larger or higher quality territories (van Schaik and van Hooff 1983; Dunbar 1988), improved access to reproductive opportunities (Snyder-Mackler et al. 2012; Roberts and Cords 2013), more efficient information transfer (van Schaik 1983; van Schaik and Burkart 2011; Lusseau, 2003), and reduced risk of predation or other attack (Janson and Goldsmith 1995; Hill and Lee 1998). Sociality also comes at a cost, for example within group competition (Cheney et al. 2004; Harris et al. 2010), or increased risk of disease transmission (Nunn et al. 2000; Creel, Schuette and Christianson, 2014). Depending on the pressures experienced by each species or group, social animals incorporate different types of social organization allowing them to maximise the benefits of social living for individual fitness (Couzin 2006; Kummer 1968; Aureli et al. 2008). The cognitive limitations of maintaining many social relationships (Sueur et al, 2011) along with limited resources in a given habitat, regulate the size and organization of group-living animals (Johnson et al. 2001). When group size increases past a threshold, intragroup competition over resources outweighs the benefits of group-living and groups may go through permanent fission events, creating two or more separate units (Nsubuga et al. 2008; Sueur and Maire, 2014). In species where kinship facilitates social relationships, closely related individuals tend to join the same group after splitting, maximising future reproductive fitness (Clutton-Brock 2016; Lefebvre et al. 2003; Nsubuga et al. 2008).

The social organisation of animal societies describes the number of individuals (e.g., 1: solitary, 2: pair-living, ≥3: group-living), demographics (e.g., number of adults, immatures; males, females), and degree of cohesion within a particular group (Kappeler and van Schaik, 2002). In addition to variation in social organisation across widespread taxa (e.g., Nishida 1968; Amici et al. 2008; Bowler et al. 2012; Patzelt et al. 2014; Forcina et al. 2019), some species also show intraspecific variation in social organisation (Caribbean cleaning goby, *E. prochilo*: Mazzei et al. 2021; sweat bee, *Halictus scabiosae*: Ulrich et al 2009; across primates: Kamilar & Baden 2014). Variation may be pronounced, with groups of the same species living in systems ranging from solitary or pair-living to group-living (Schradin et al 2018); while in other cases all individuals are group-living, but these groups vary in their size, sex ratio, or cohesion (e.g., *Physter macrocephalus*: Whitehead et al. 2012; *Elephas*: de Silva and Wittemyer, 2012).

In larger groups variation between individuals may lead to the adoption of different behavioural strategies or roles, for example, when mating competition is high, some males may adopt satellite or peripheral roles to avoid high-risk competition with other more dominant or central males (Berec and Bajgar, 2011; Howard 1978; Balmer et al. 2019; Setchell, 2003). Female chimpanzees may occupy central or peripheral positions within their group, with peripheral females able to avoid feeding competition from larger parties (Williams et al. 2002). In black tufted-ear marmosets (*Callithrix kuhli*), as in many callitrichids, cooperative-breeding groups are formed that contain both breeding individuals and helpers who offer additional parental care to young without breeding (Schaffner and French, 1997).

Group size itself may also be a determining factor in the social organization of some species. Larger groups are more likely to exhibit intermediate subgrouping patterns (Kappeler 2019; Grueter et al, 2012; Aureli et al. 2008; Couzin and Laidre, 2009) due to potential upper limits on the energy available to maintain social relationships (Sueur et al. 2011). Fission-fusion dynamics, which describe the spatial cohesion and membership of temporary parties formed by a subset of individuals from within the larger group, provide a range of solutions for individual flexibility in social living (Aureli et al, 2008). In ‘atomistic’ fission-fusion systems (also referred to as *higher* fission-fusion (Aureli et al. 2008)) individuals vary the number and composition of their immediate associates (their subgroup) – joining or splitting from other individuals (from within their social group or community) fluidly across the day and allowing individuals to range independently while being members of a larger social unit (Aureli et al. 2008; Grueter and Zinner, 2004: chimpanzees (*Pan troglodytes*, Lehmann and Boesch, 2004), spider monkeys (*Ateles sp*. Symington, 1990), spotted hyenas (*Crocuta crocuta*, Holekamp et al. 2012), humpback dolphins (*Sousa sahulensis*, Hunt et al. 2019), big brown bats (*Eptesicus fuscus*, Metheny et al. 2008), and blue, marsh, and great tits (*Paridae sp*. Aplin et al. 2014)). Multilevel social organisation describes nested fission-fusion social dynamics (Aureli et al. 2008; Grueter et al. 2004; Grueter et al. 2012; Grueter et al. 2020), with fissioning and fusing taking place not on the level of the individual, but on core subunits that maintain reliable, stable cohesion, and static membership while nested within a larger social group with two or more levels (Grueter et al. 2020; Grueter and Zinner., 2004; for example in: geladas (*Theropithecus gelada*, Snyder-Mackler, Beehner and Bergman, 2014), African elephants (*Loxodonta africana*, de Silva and Wittemyer, 2012), plains zebras (*Equus quagga*, Grueter et al. 2017; Rubenstein and Hack, 2004), hamadryas baboons (*Papio hamadryas*, Schreier and Swedell, 2009), and sperm whales (*Physeter macrocephalus*, Rendell and Whitehead, 2003)). The presence of fission-fusion dynamics already implies some level of flexibility in social organization; however, the drivers for intraspecific variation may vary across species and timescales.

The long-term combination of multilevel and atomistic fission-fusion sociality has been suggested to be a point of distinction in the evolution of human social organisation, enabling humans to cooperate flexibly across very large numbers of unrelated people (Grueter et al. 2012, 2020). Our closest living relatives, chimpanzees and bonobos, both employ atomistic fission-fusion sociality (Nishida 1968; Lehmann and Boesch, 2004; Kuroda 1979; Surbeck et al. 2017). However, before the permanent fissioning of two chimpanzee communities (Reddy, 2020; Sandel and Watts, 2021; Feldblum et al. 2018), two studies found that, in additional to their atomistic fission-fusion dynamics, males were further divided into two ‘cliques’ (or subgroups) that were either stable over a two-year period (Ngogo; Amsler and Mitani, 2003) or formed rapidly over the two years prior to fissioning (Gombe; Feldblum et al. 2018). Large community size may also contribute to the presence of four female cliques in Ngogo, however, the competitive pressures driving these cliques may differ between sexes (Wakefield 2008). The integration of an additional level of social organisation in the Ngogo and Gombe chimpanzees (Mitani and Amsler 2003; Feldblum et al. 2018) may have been a short-term strategy to offset the costs of living in a large community or in a community with a heavily male biased sex ratio (Ngogo = 1:1.15; Mitani, 2002; Gombe ∼1:1, operational sex ratio 5:1 before split; Feldblum et al. 2018). However, these are the only groups of chimpanzees in which grouping patterns beyond atomistic fission-fusion have been reported to date. To strengthen the case for a relationship between group size and the use of additional levels of structure with atomistic fission-fusion sociality we need to further explore variation in the social organization of chimpanzee communities of different sizes and compositions.

Chimpanzees are highly social primates, living in communities (or ‘unit groups’) of typically 50-80 individuals (Nishida 1968; Goodall 1986; Nishida et al. 1990). A single temporary subgroup or party, rarely, if ever, includes all individuals from one community, with party size and composition varying substantially throughout the day (Nishida 1968; Goodall 1983). General community cohesion appears to be impacted by resource availability (Matthews et al. 2021), inter-community competition, predation risk (Lehmann and Boesch 2005; Boesch 1991), total community size, and the relative number of adult males, with smaller communities and those with smaller numbers of males exhibiting higher cohesion (Lehmann and Boesch 2004). Males are philopatric and highly gregarious (Goodall 1986), and while kinship plays a role in male-male association (Sandel et al. 2020) most relationships occur between non-kin (Langergraber et al. 2007). Intercommunity interactions are typically hostile and can be lethal (Mitani et al. 2010; Wilson et al. 2014); and the high levels of tolerance and coalitionary support required to maintain and defend territorial space are often maintain through non-kin social bonds (Mitani 2009; Gilby et al. 2013; Samuni et al. 2021).

A few cases of very large chimpanzee communities (100-200 individuals) have been reported (Mitani and Amsler 2003; Samuni et al. 2014). Communities with larger numbers of males are able to maintain larger territories (Lemoine et al. 2020), which in turn sustains larger numbers of females and increased reproductive opportunities (Mitani et al. 2010; Lemoine et al. 2020). However, a larger number of males also increases within-group competition. While one strategy to mitigate within-group competition is via post-conflict management mechanisms (Preis et al., 2018; Kutsukake and Castle, 2004), time-allocation constraints likely hamper the effective management of social relationships in large groups (Lehmann et al. 2007; Sueur et al. 2011). An additional level of social organisation may, up to a point (c.f. Sandel and Reddy 2021), mitigate increased male-male competition and allow for the formation of ‘super-communities’ with the potential to outcompete more typically sized neighbouring groups (Mitani and Amsler, 2003).

We test the hypothesis that atomistic fission-fusion chimpanzee communities with many males should incorporate additional levels of male social organisation and strategies by exploring patterns of social association in a large group of chimpanzees, the Waibira community of the Budongo Forest, Uganda, who have an estimated 120 individuals and 24-31 males that travel independently of their mothers. We compare the social organisation of this community to that of their neighbouring community, Sonso, who present a more typical East African chimpanzee demography with around 65-70 individuals, and 10-13 males. We contrast the level of cohesion and subgrouping within both networks, and explore the stability of subgrouping, and the impact of social grouping on ranging behaviour in Waibira males.

## METHODS

### Study Site and Subjects

Observational data from two habituated chimpanzee communities: Sonso and Waibira, were collected by long-term field assistants between October 2015 and October 2019 in the Budongo Forest Reserve, Uganda (1°35’ -1°55’N, 31°18’-31°42’E). The Sonso community has been observed since 1990, while observations of the Waibira community started in 2011. During the study period the Waibira community was composed of between 83 and 94 identified individuals (including infants and juveniles: 0 - 10 years) with between 24 and 31 independent males (travelling independently of their mother and observed in parties without their mother). There are also several peripheral females with dependent offspring in the Waibira community who have not been formally identified, increasing the estimate of community size to ∼120 individuals. Across the four years of observations, we had 34 individually identified independent males in the Waibira community. Three of these were discarded from analysis because they disappeared/died more than six months before the end of the study period and a further nine were discarded because they reached independence more than six months after the start of the study period, leaving 22 males in the Waibira dataset. The Sonso community included between 64 and 68 identified individuals (including infants and juveniles) with between 10 and 13 independent males. Across the four years of observations, we had 14 individually identified males in the Sonso community. Three of these were discarded from the analysis because they disappeared/died more than six months before the end of the study period, leaving 11 males in the Sonso dataset.

We include all males who make travel decisions independently from those of their mother (independent males), as these individuals influence the patterns of male social organisation of the community through their grouping decisions. In addition, by independence, males begin exhibiting adult behaviour through mating (Crockford et al. 2020; Constable et al. 2001; lower-bound for paternity in Budongo is 9-years), engaging in boundary patrols (Mitani and Watts, 2001), and forming strong social bonds with adult males (Sandel et al. 2020). The transition from subadult to adult is fluid, variable, and not reliably predicted by age alone (e.g., Reynolds, 2005; Goodall, 1986), by including all independent males we also avoid making potentially arbitrary decisions about the minimum age for males to be considered adults.

### Party Composition Data

Party composition data were extracted from the Budongo long-term dataset for the period between October 2015 and October 2019. Party composition scans were recorded every 15 minutes during daily focal follows (Altmann 1974) by trained field assistants in both communities. A party was defined as a subset of independent individuals from the community exhibiting coordinated behaviour within a rough 35-meter radius of the group centre (Newton-Fisher 1997). During the party composition scans, location of the party and the identity of all independent individuals were recorded. The location corresponds to a ‘block’ within a grid system of North to South and East to West trails, which marks out the majority of the communities’ ranges.

### Data manipulation

There are typically between two and four field assistants collecting behavioural data from each chimpanzee community every day. Due to the fission-fusion dynamics of chimpanzee communities, multiple field assistants may record the same party at the same time if the individuals they are following that day join the same party. We deleted duplicate parties (parties recorded in the same block, at the same time, with an overlap of individuals) by removing the smaller party from the dataset. If duplicate parties were the same size, the party recorded by the more experienced field assistant was retained.

As chimpanzees can remain in one place (e.g., a feeding tree) for more than 15 minutes, parties collected within 15-minutes of each other may not represent independent samples. To account for the possible influence of non-independence of parties on our results we conducted our analyses twice - once using an individual randomisation to construct null models based on the Farine (2017) permutation method, and once using a subset randomisation method from Surbeck et al. (2017). The individual randomisation method controls for individual differences in number of observations (more below), whereas the subset randomisation method controls for the non-independence of consecutive parties. In the subset randomisation we randomly selected a subset of parties from the original dataset at a mean interval corresponding to the mean number of consecutive parties that dyads were seen together. For both Waibira and Sonso subset randomisation resulted in a mean interval of six scans. These subsets were used to run the analysis and null models were constructed using the same method as the individual randomisation but using the subset data.

Blocks within the grid system vary in size because the distance between trails varies, and the presence of some blocks changed over time as new trails were cut. Blocks were combined to produce a new grid system in which all blocks were 200 × 500 meters and were consistent across all the years included in ranging analyses. In doing so we were able to control for uneven sampling between blocks due to their size. Where the block was not recorded or was unclear (e.g., mistyped) these scans (N=579, 0.03%) were omitted from ranging analyses. The block locations were converted to coordinates using UTM latitude and longitude data points taken at the South-West corner of each block with a Garmin GPS device.

### Social network analysis

We constructed weighted social networks (Farine and Whitehead 2015) to represent the associations between independent males (hereafter males) within each community using the ‘igraph’ package (Csárdi 2006) in R v4.04 (R Core Team 2021) on a MacBook Pro with Catalina. Edge weights were quantified using a dyadic association index (DAI), also known as simple ratio index (Cairns and Schwager 1987), calculated as follows:

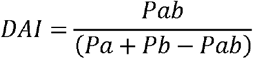

Where *Pab* is the number of time individuals ‘*a*’ and ‘*b*’ were seen in a party together, *Pa* is the number of times individual ‘*a*’ was seen in a party and *Pb* is the number of times individual ‘b’ was seen in a party.

Mantel tests were used to compare the overall association patterns of each community across years to determine if the patterns of association between individuals remained stable over time. Here, we compared association matrices created using DAI for every year with all other years (Morrison et al. 2019).

We constructed null models for these networks using the ‘asnipe’ package in RStudio (Farine 2019), following the methods described by Farine (2017) using data stream permutations of the raw data for the full four-year datasets available for each community with 10,000 permutations. We conducted four permutations in total, one for each community (Sonso and Waibira) and one for each detected subgroup (Waibira_A and Waibira_B). The permutations randomly swapped individuals between parties, thereby keeping the party size and the number of times individuals were observed constant while changing party composition.

We quantified the organisation of these networks using the following measures: Transitivity, mean Strength and Modularity. Transitivity was used to measure the probability of triangular connections (three nodes being connected to each other). This measure represents the cohesiveness of the network (Barabási and Albert 2002). The mean Strength of nodes within the network was calculated by summing the edge weights of each node (Farine and Whitehead 2015) and averaging across all nodes in the network. Individual Strength measures the sum of individual edge weights providing a measure of the overall quality of an individual’s associations. Therefore, mean Strength may be used to infer the relative degree of dyadic association within chimpanzee networks (e.g., McCarthy et al. 2019). Finally, we identified subgroups within each community using Louvains clustering algorithm (Blondel et al. 2008; Morrison et al. 2019) and used Modularity to quantify how well the networks divided into these subgroups. Modularity measures the likelihood of edges being present within compared to between subgroups. We compared these measures to the null models to check if they represented non-random characteristics of the networks (Morrison et al. 2019). The resulting network measures were considered non-random if the observed network measures were higher or lower than more than 95% of the values generated by the null model using the following formulae:

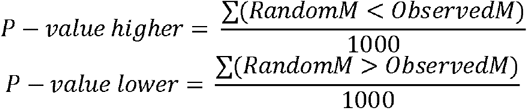

Where RandomM is the distribution of values in network measures obtained from the null model and ObservedM is the observed values of the true network measures (Farine 2017). By using three network measures we were able to address multiple characteristics of the community’s social organisation that could vary between communities.

To verify whether our findings were an outcome of the inclusion of subadults in our dataset, we replicated the first section of our analysis (calculations of network measures and differences in network structure between communities) using only adult individuals (aged 16-years or older, Reynolds, 2005) creating networks of seven and 15 adults in Sonso and Waibira respectively. These adult-only analyses are provided in the Supplementary Materials.

### Does subgroup membership predict individual strength?

If subgroups were found in a community, we used a linear model to test the relationship between individual Strength and subgroup membership. Next, we compared the coefficient from this model to coefficient estimates generated from the null model. If the observed coefficient was smaller or greater than 95% of the of the coefficients generated from the null models, we considered this effect stronger than expected by chance (Farine 2017). P-values were calculated using the same method as described above. We could only consider the impact of individual Strength on group membership because this was the only individual-level measure we calculated. This analysis allows us to identify the presence (or absence) of behavioural differences between individuals in different subgroups with regards to their association patterns. If the null hypothesis is rejected, then subgroups are characterized by differences in individual Strength. Differences in individual Strength may reveal different behavioural strategies employed by individuals in different subgroups, in this case, with regards to the gregariousness of individuals.

### Comparing network structure between communities

We used the dissimilarity measure proposed by Schieber and colleagues (2017) to compare the overall structure of networks between communities and subgroups. This method allows us to compare the distribution of edges (associations) in relation to nodes (individuals) across two networks while controlling for difference in network size. Outputs from this computation range from zero to one, with value close to zero implying high *similarity* between networks and values approaching one indicating greater *dissimilarity* between networks. This method identifies topological differences between networks of the same and different sizes by comparing three Probability Distribution Functions (PDFs) that define the ways in which nodes are connected to each other within the network. The PDFs include the network distance distribution (which defines global topological properties of the graph by comparing average connectivity of nodes), the network node dispersion (which describes the degree of heterogeneity of node connectivity), and the alpha-centralities of the networks (which describes the direct and indirect way in which nodes are connected in the network and captures the effect of disconnected nodes). Detailed information about the way the PDFs are calculated can be found in Schieber et al. (2017). One drawback of this method is that it cannot be used for weighted networks. To overcome this limitation, we converted our networks into binary unweighted networks, retaining only the edges that represented associations that occurred above chance while including all nodes (individuals) from the community (Joudaki et al. 2012). As a result, some nodes were not associated with any edges and not connected to any other nodes (because those individuals did not share any above chance associations). To calculate the chance level of association we calculated the mean edge weight of 10,000 random networks constructed using the permutation method described above. Edges with weights below or above chance were set to 0 or 1, respectively.

### Home range analysis

We used the ‘adehabitatHR’ package (Calenge 2020) in RStudios to calculate the size of each community’s home-range using the top 99% of the most frequented locations where individuals were observed. Where modules were identified within communities, we used the ‘kernaloverlap’ function to calculate the proportion of home-range overlap between each possible male dyad in the community. We carried out this analysis using 5%, 25%, 50%, 75% and 95% of the individual estimated home ranges. Finally, we assessed whether the home range overlap of dyads within the same subgroup was larger than that of dyads from different subgroups using an unpaired permutation Student’s t-test with 1000 permutations. For this test we used the ‘RVAideMemoire’ package in RStudios (Hervé 2020).

From all parties recorded in the party composition scans, we extracted parties that included individuals from each subgroup. We then used the ‘heatmap.2’ function in RStudios to map the proportion of scans where parties with males from each subgroup were recorded in each block location within the community home range. We created separate heat maps for each subgroup. These figures were used to identify any differences in ranging patterns between subgroups. To control for potential daily variation in the number of parties recorded, and number of consecutive parties recorded in a given block we used only the first party recorded each day.

### Stability of subgroup membership

Where subgroups could be identified, we investigated whether subgroup membership was stable over time. We split our dataset by year (N = 4) and used Louvain’s clustering algorithm to determine the presence of subgroups in each of the four years separately. We then used the Levenshtein Distance to measure the absolute difference in membership between a specific subgroup across each pair of consecutive years. Levenshtein distance measures the least expensive path between two sets of strings (in this case individuals IDs within a subgroup) by counting the number of insertions, deletions or substitutions required to get from one string to the next (Kessler 1995). For example, the distance between a subgroup including BEN, LAF, MUG, TAL to a subgroup containing BEN, LAF, LAN, TAL, MAS is two (one substitution between MUG and LAN and one insertion of MAS).

## RESULTS

### Overall network structure

After deleting duplicates, we retained 41,560 party composition scans across 1,248 days from the Sonso community and 19,117 scans across 1027 days from the Waibira community. Individual males in the Sonso community were present in a mean of 12,053 scans across the four-year study period (range: 10,104-15,051, SD = 1,715.12), while males from the Waibira community were observed in a mean of 2,540 scans per individual (range: 68-6,134, SD = 1,963.12). The mean number of males in parties that include at least one male was 4.66 in Sonso (range = 1-14, SD = 3.62) and 4.82 in Waibira (range = 1-20, SD = 3.81). At the start of the study period males in Sonso were between nine and 23 years old (mean = 16.9 years) and males in Waibira were between 10 and 39 years old (mean = 20.73 years). Mantel tests revealed that male-male dyadic associations in Waibira and Sonso remained stable over the four-year study period (Supplementary Material Table S1; Figure S1). The Sonso home range (99% Kernel) was approximately 5.33km^2^ across the full study period, while the Waibira home range was approximately 10.28km^2^.

Analyses from the individual randomisation and subset randomisation revealed quantitively similar findings. We report here the results from the individual randomisation, which retained a larger dataset. Results from the subset randomisation can be found in the supplementary materials (Supplementary Materials Tables S2-S5, Figure S2-S4). Furthermore, results did not change when including only adult individuals (Supplementary materials Table S6-S7, Figure S5).

Transitivity of both networks was 1, suggesting that over this study period all male dyads in both communities were observed in the same party at least once. This measure was not significantly different from the null models (Figure 1, P = 1). Such a high transitivity confirms that both communities represent two distinct and coherent groups in which, over a four-year period, all male dyads in both communities were observed in the same party at least once.

**Fig 1.**
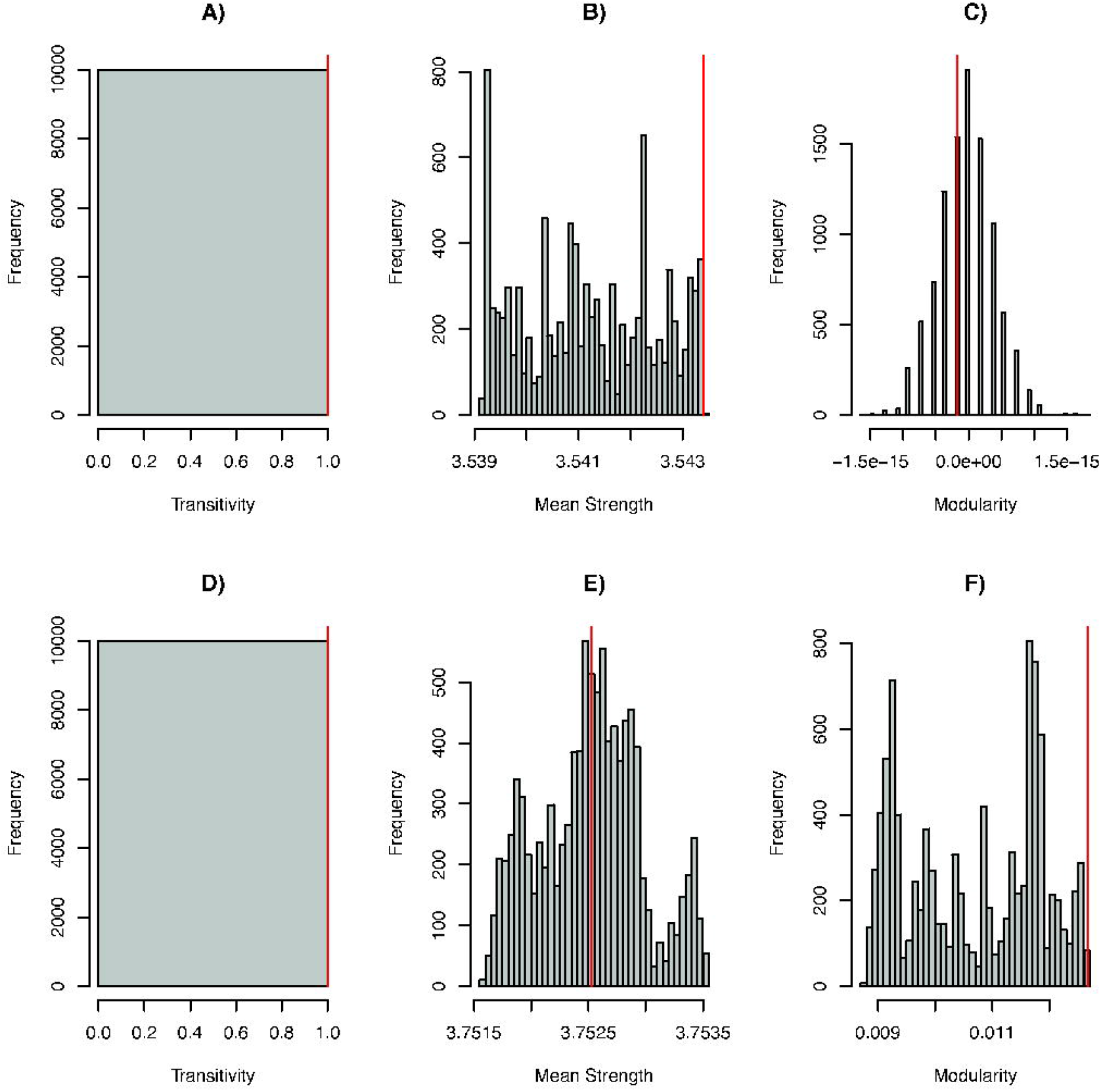
Histogram distribution of a) Sonso Transitivity, b) Sonso mean Strength, c) Sonso Modularity, d) Waibira Transitivity, e) Waibira mean Strength and f) Waibira Modularity measures from 10 000 data stream permutations of the Sonso and Waibira male social networks. Red lines indicate measures from networks created from original party composition data

Individual Strength (sum of relationship edge-weights) ranged between 2.6 and 4.24 (mean = 3.54, SD = 0.42) in the Sonso community and between 1.14 and 5.09 (mean = 3.75, SD = 1.08) in the Waibira community. The mean Strength across the entire four-year dataset was greater than expected by chance for individuals in the Sonso community (Figure 1, P < 0.001), but not in Waibira, (Figure 1, P(greater) = 0.51 and P(smaller) = 0.491).

Using Louvain clustering algorithm, we identified one module in the Sonso community encompassing all independent males (Figure 2); and Modularity in the Sonso community almost negligible (−1.82^-16^) and not different from chance (Figure 1, P(greater) = 0.564 and P(smaller) = 0.436 respectively). The small modularity observed in Sonso suggests no clustering was detected in the Sonso community. Two modules were identified in the Waibira community, separating the community into subgroups of nine (Waibira_A) and 13 (Waibira_B) individuals (Figure 2); Modularity in the Waibira community was greater than expected by chance (Figure 1; P = 0.002). However, Modularity in the Waibira community remained low (0.013) suggesting that this pattern of subgrouping was not robust (Newman, 2006). Given the high atomistic fission-fusion dynamics of this community and the finding that all individuals associated with each other at least once within the study period (Transitivity =1), low modularities are expected. Visual inspection of the Waibira network (Figure 2) suggests that while the Waibira_A module appears to represent a cohesive subgroup of males who associate preferentially with each other, the Waibira_B module does not seem to form a cohesive subgroup, but instead represents satellite males who did not associate preferentially with specific individuals. Finally, modularity values in Waibira varied across years – suggesting some flexibility in subgrouping between male chimpanzees, as described below (Figure 5; Table 3; Supplementary Materials Table S9). However, modularity in Sonso remained around zero across all four years (Table 1) supporting the finding that Sonso males do not divide into subgroups.

**Table 1.**
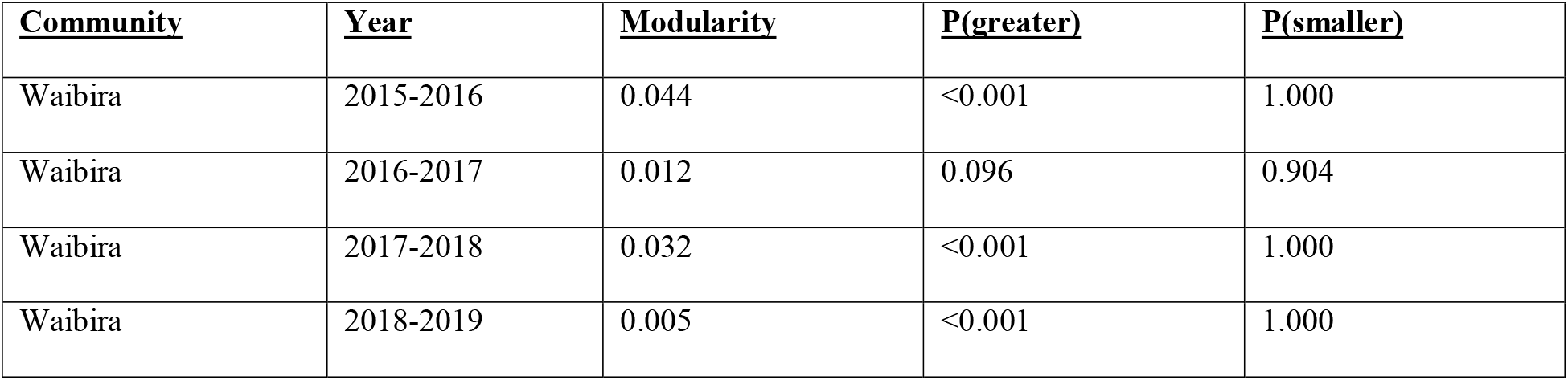
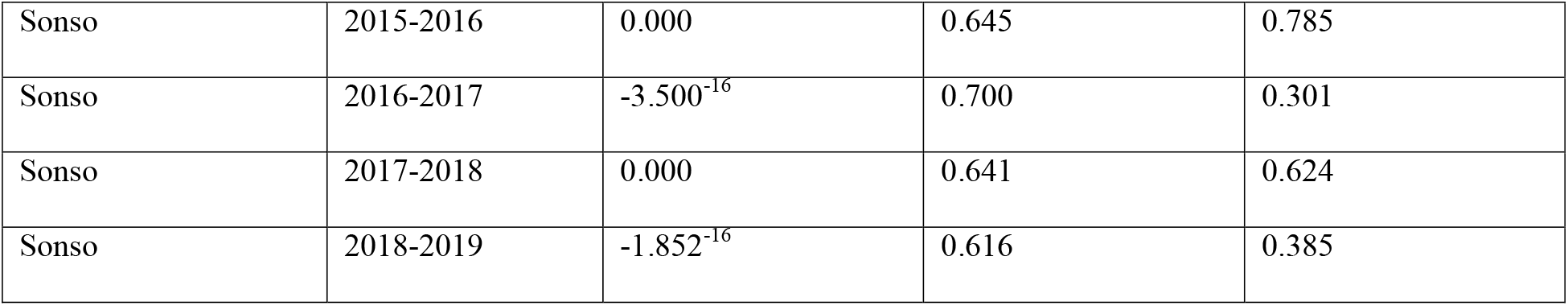
Modularity of Waibira and Sonso community across four-year study period. Including the P-value generated by comparison with null models (10,000 data stream permutations).

| <u>Community</u> | <u>Year</u> | <u>Modularity</u> | <u>P(greater)</u> | <u>P(smaller)</u> |
| --- | --- | --- | --- | --- |
| Waibira | 2015-2016 | 0.044 | <0.001 | 1.000 |
| Waibira | 2016-2017 | 0.012 | 0.096 | 0.904 |
| Waibira | 2017-2018 | 0.032 | <0.001 | 1.000 |
| Waibira | 2018-2019 | 0.005 | <0.001 | 1.000 |
| Sonso | 2015-2016 | 0.000 | 0.645 | 0.785 |
| Sonso | 2016-2017 | -3.500<sup>-16</sup> | 0.700 | 0.301 |
| Sonso | 2017-2018 | 0.000 | 0.641 | 0.624 |
| Sonso | 2018-2019 | -1.852<sup>-16</sup> | 0.616 | 0.385 |

**Fig 2.**
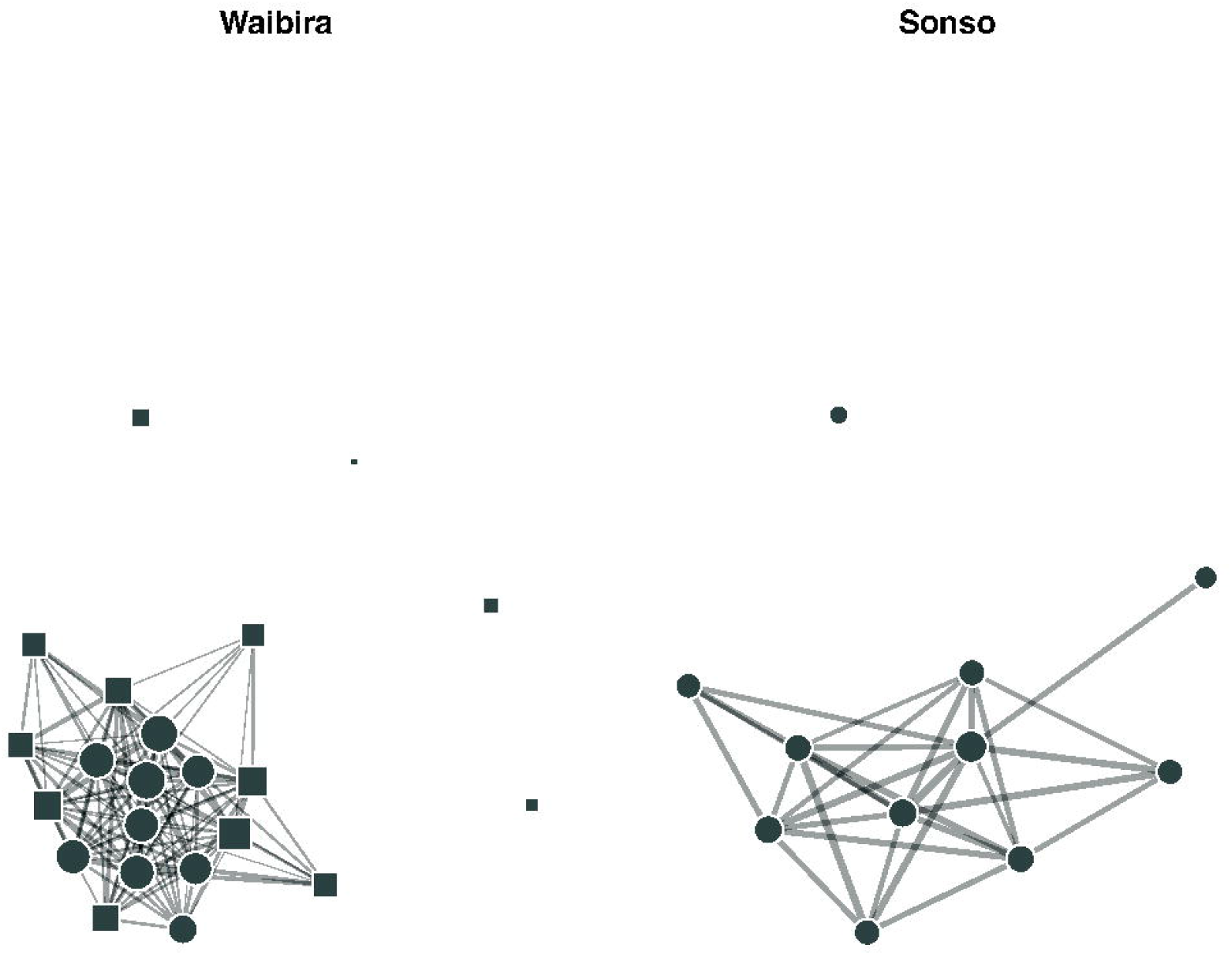
Sociogram representing the Waibira (left) and Sonso (right) male social networks. Edge weights represented the dyadic association index between two individuals multiplied by seven (for clearer visualisation). Only edges with a value higher than chance from 10 000 random permutations (Sonso = 0.354, Waibira = 0.18) are present in the network. Node shapes within each network indicate subgroup membership. Node area represents individual Strength multiplied by five (for clearer visualisation), so that larger nodes have higher individual Strengths

### Do males in Waibira_A and Waibira_B exhibit different individual Strength?

We tested whether membership in Waibira_A or Waibira_B was related to the quality of association Strengths individuals had with all other individuals in the community. A linear model revealed that individuals in Waibira_A had stronger individual Strengths as compared to individuals in Waibira_B (Figure 3, linear model: F = 20.61, DF1,20, R^2^ = 0.48). This finding was also different than expected from the null model (P < 0.001) and suggests that males in Waibira_A formed stronger associations with other males as compared to males in Waibira_B.

**Fig 3.**
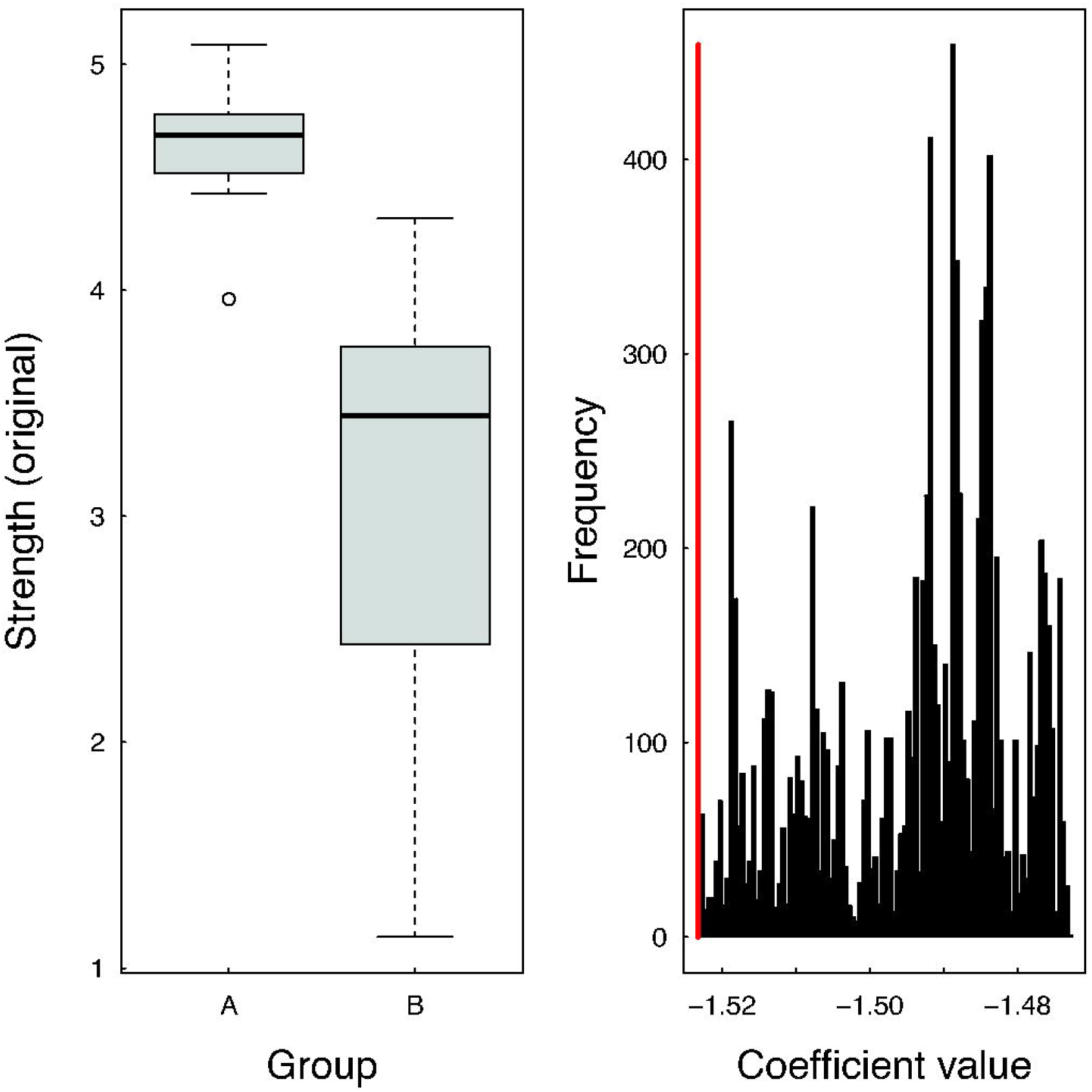
Illustration of model results testing the relationship between individual Strength and subgroup membership in the Waibira community. a) boxplot illustration of the range of individual Strength in each group with thick lines illustrating the mean Strength in the community, box limits are the lower (25%) and upper (75%) confidence intervals, and the range shows the most extreme (highest/lowest) Strengths that are no more than the range multiplied by the interquartile range. b) the distribution of model coefficients from the null model; the red line indicates the coefficient from the model testing the original data

To get a better understanding of this difference in gregariousness, we tested whether males in Waibira_A and Waibira_B differed in other aspects that may impact their gregariousness: their age and rank. At the start of the study period males in Waibira_A showed an older mean age (mean = 23 years, range = 12-39) than males in Waibira_B (mean = 19, range = 10-34). At the end of the study period males in Waibira_A showed a higher mean rank (mean rank = 8.77, range = 1-21, Supplementary Materials Figure S6) as compared to males in Waibira_B (mean rank = 13.38, range = 4-22; see Supplementary Materials for dominance rank assessment, Figure S6); however, the range of ages and ranks represented in both groups showed almost complete overlap.

### Dissimilarity between network structure of Waibira and Sonso

Using the *dissimilarity* method, we compared the structure of the Sonso and Waibira networks independently of community size to understand if the differences detected between the communities were related to the social organisation of the two groups. As our clustering algorithm detected two modules in the Waibira community we also compared the Sonso community network to the networks of each Waibira module (Waibira_A and Waibira_B) separately. By converting our networks into binary, unweighted networks, we retained the following number of edges (= 1) in each network: Sonso = 29/55, Waibira = 119/231, Waibira_A = 16/36, Waibira_B = 18/78. This analysis revealed the strongest similarity between Sonso (n=11) and Waibira_A (n=9), and the weakest similarity between Sonso and Waibira_B (n=13).

**Table 2:**
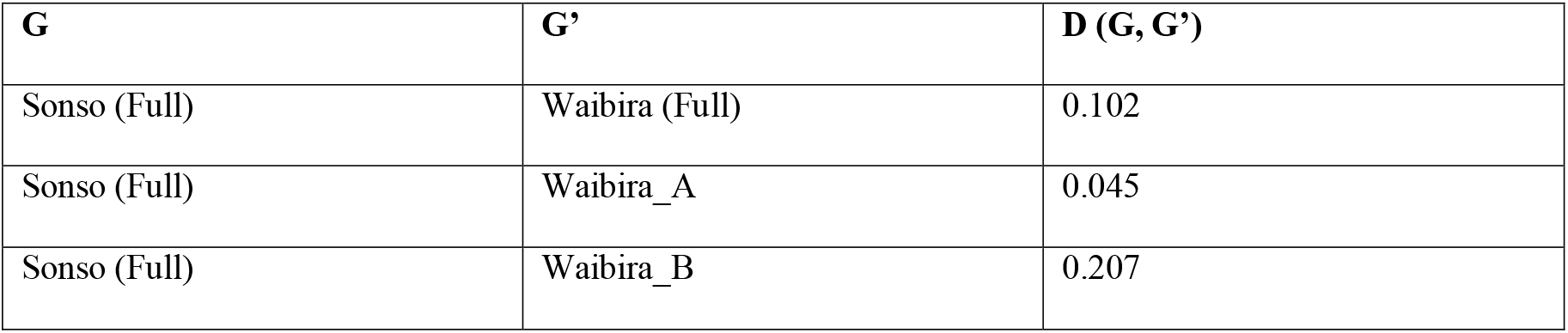
D dissimilarity values between communities and between Sonso and Waibira subgroups. G is the community/subgroup to be compare with G’ (another community/subgroup)

### Do individuals from Waibira_A and Waibira_B share an overlapping home-range?

Out of 15,646 parties recorded within known blocks in the Waibira grid system with at least one male present, individuals from Waibira_A were observed in 13,463 and individuals from Waibira_B were observed in 12,051 scans. 3,595 scans included only individuals from Waibira_A, 2,183 included only individuals from Waibira_B, and 9,868 included individuals from both subgroups. We found no difference in the ranging habits of the two Waibira subgroups. Males were not more likely to have overlapping home-ranges with individuals within their own subgroup as compared to those in the other subgroup when taking the first observation of the day (Supplementary Materials Table S8; Figure 4). This finding did not change when including all observed parties recorded throughout the day (Supplementary Materials Table S9; Figure S7)

**Fig 4.**
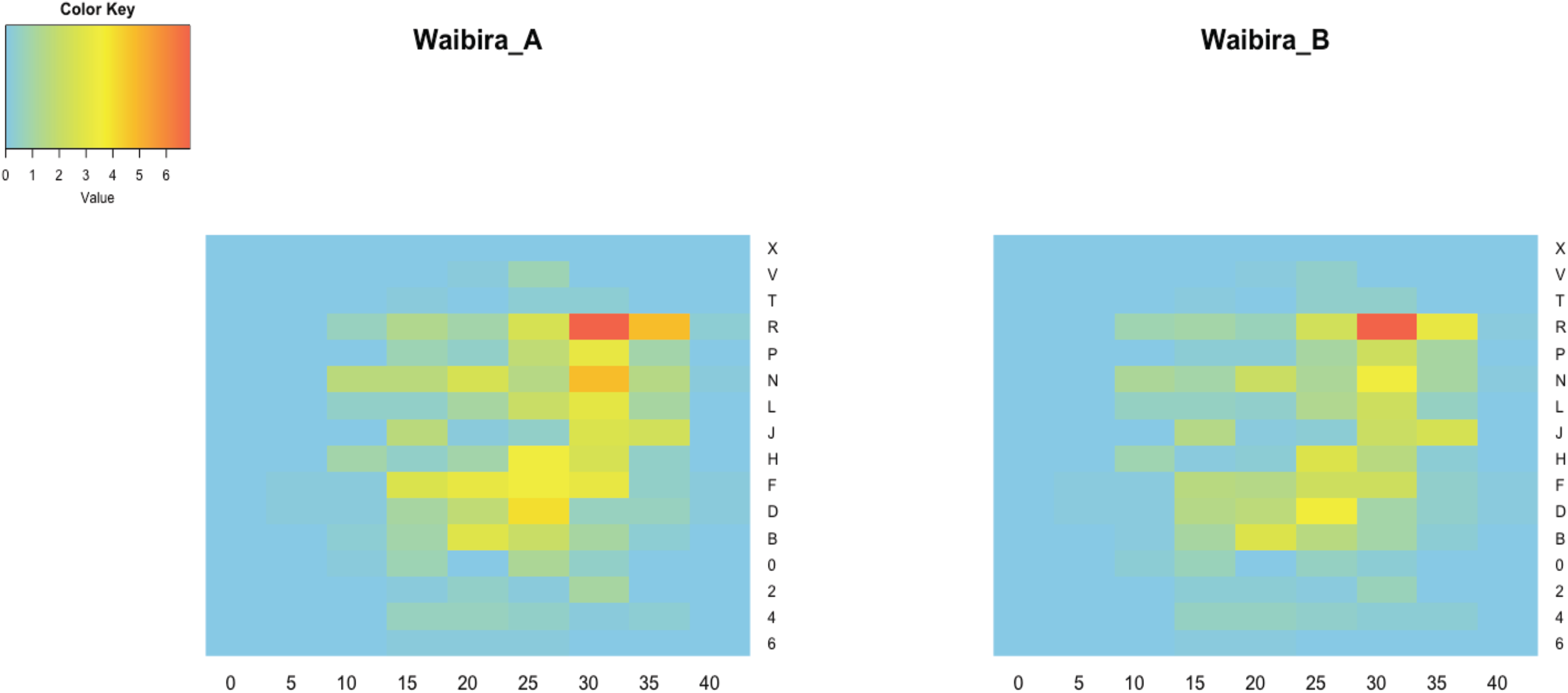
Heat maps showing the ranging patterns of Waibira_A and Waibira_B from the Waibira community. These maps take only the first observation of each day and includes all parties in which one or more individuals from the group were present. Colour key indicates the proportion scans where individuals from each group were observed in each block with red indicating highest proportion time in the block and blue indicating no time spent in the block

### Stability in Waibira subgroup membership

While the network measures described above allow us to assess the presence of subgrouping within Waibira, they do not assess the stability of the membership of these groups. Across the four years, the number of subgroups identified, the membership, and the mean Strength of these subgroups changed (Figure 5; Supplementary Material Table S9).

**Fig 5.**
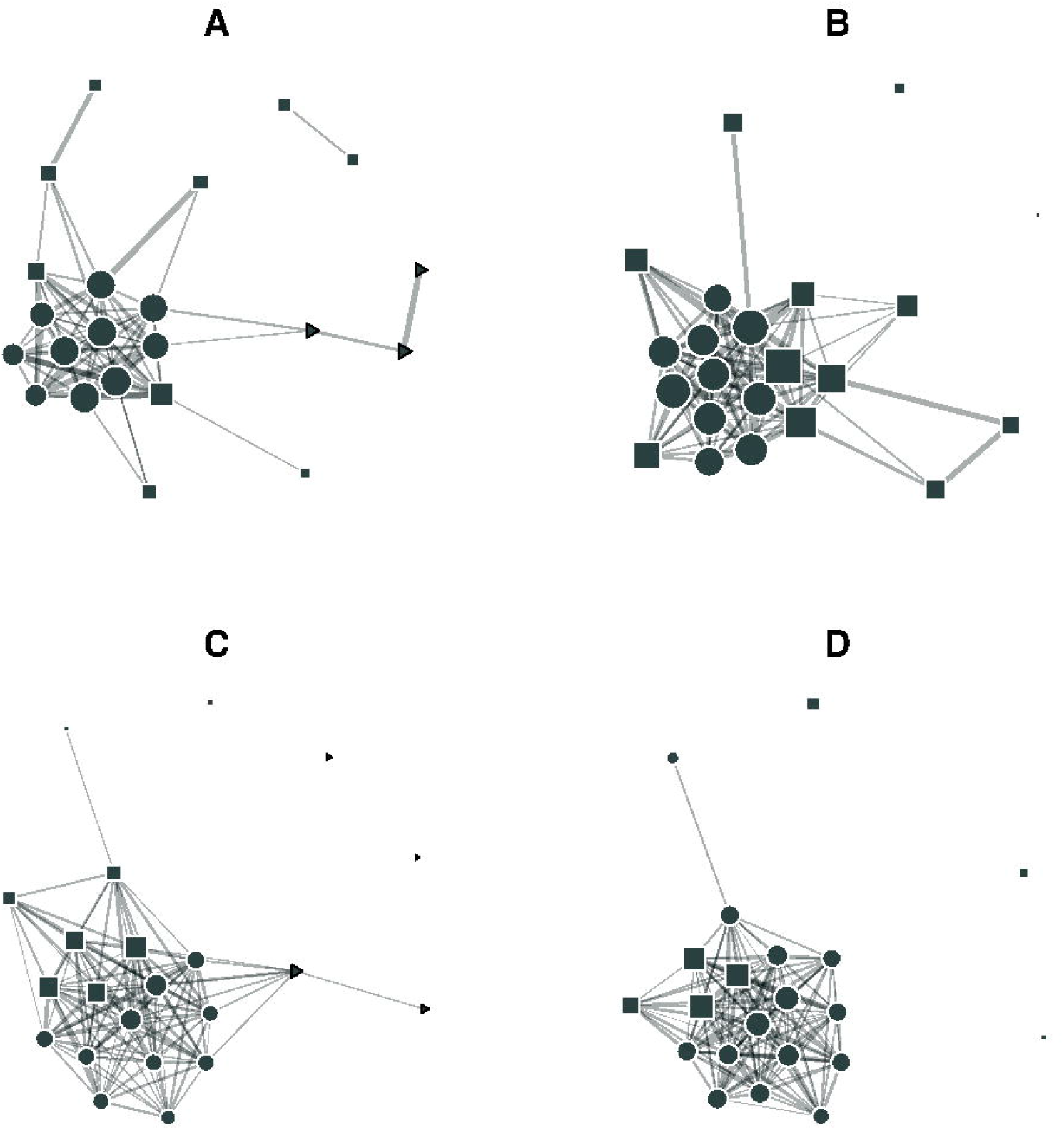
Sociograms highlighting subgroups identified through Louvain’s clustering algorithm between males in the Waibira community for each of the four years of study: A) 2015-2016, B)2016-2017, C) 2017-2018, D) 2018-2019. Only edges with a value higher than the community mean (2015-2016 = 0.169, 2016-2017 = 0.22, 2017-2018 = 0.123, 2018-2019 = 0.146) are present in the network. Node shapes within each network indicate subgroup membership. Node area represents individual Strength multiplied by seven (for clearer visualisation), so that larger nodes have higher individual Strengths. Circular nodes always belong to ‘Waibira_A’ or ‘Waibira_A1’: the subgroup with highest mean Strength

The Waibira_A subgroup, which contained individuals with relatively high mean Strength, was present across most years. We found very high membership stability in Waibira_A over the first two study years (2015/2016 and 2016/2017). In the third study year (2017/2018), this subgroup appeared to split into two smaller subgroups (Waibira_A1 and Waibira_A2), one of which (Waibira_A1) included new members who had moved over from the Waibira_B subgroup. In the final year (2018/2019) the two Waibira_A subgroups from the previous year (A1 and A2) appear to merge to form one supergroup still including most of the Waibira_A individuals present across all years, as well as others from Waibira_B who had moved in (Figure 5 and Table S9).

## DISCUSSION

Following findings from previous research on the particularly large Ngogo chimpanzee community (Amsler and Mitani, 2003), we tested the hypothesis that chimpanzee communities with many males exhibit modular social organisation in another unusually large community of chimpanzees (the Waibira community). We predicted that, while Waibira should exhibit modular social organisation with multiple subgroups, a ‘typically’ sized group (the Sonso community) will not. We did not find evidence for distinct subgrouping or robust modularity in either community; however, we did confirm the presence of flexible social organisation – including one cohesive module composed of a subgroup of males – existing within the atomistic fission-fusion society of the larger Waibira community of chimpanzees. We show new diversity in male chimpanzee social organization and social strategies within the larger community – the presence of satellite males surrounding a cohesive male subgroup - not detected in the smaller community or indeed in previous studies of large communities.

**Table 2.**
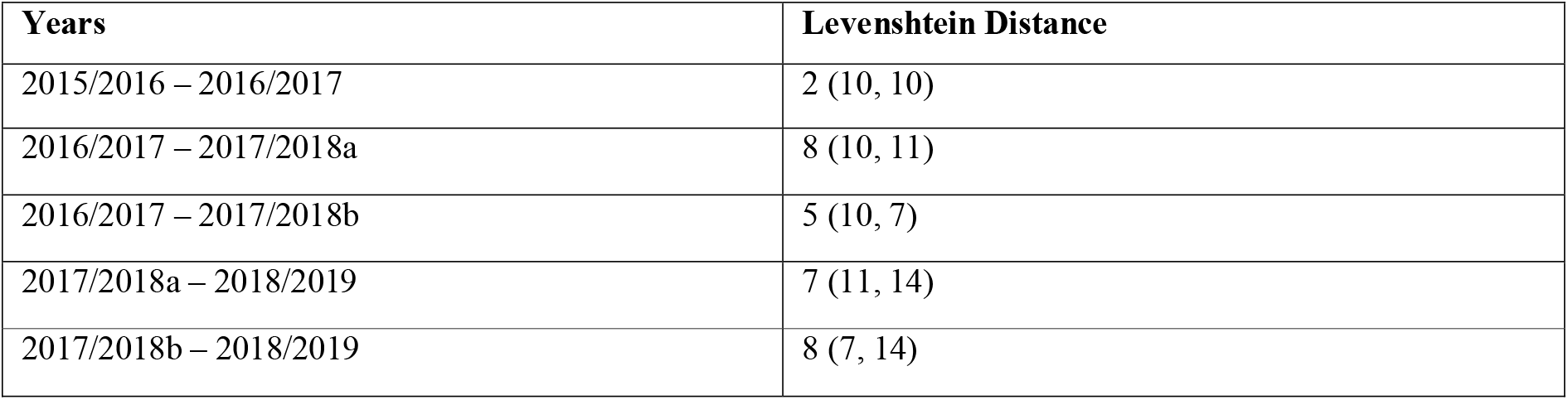
Levenshtein Distance between Waibira_A subgroups across each pair of consecutive study years All comparisons are made for the Waibira_A subgroup identified across the four-year study period. Waibira_A was defined as the subgroup with the highest mean Strength. During the 3^rd^ study year two subgroups were identified with similarly high mean Strength so we named these Waibira_A1 (2017/2018a) and Waibira_A2 (2017/2018b). In this table we report the Levenshtein Distance between two Waibira_A subgroups of consecutive years and the number of individuals in the subgroup for each year in parentheses. E.g., between the first two study years the Levenshtein distance between the Waibira_A subgroup was 1 and the subgroup size for both years was 10 individuals, we report this as 1 (10, 10)

Waibira and Sonso form two distinct, intact chimpanzee communities in which all males range in parties with all other males from that community at some time. Both communities show ‘atomistic’ fission-fusion dynamics and, despite differences in community size and number of independent males, parties in both Waibira and Sonso tended to contain the same number of males. However, while Sonso males employ the atomistic fission-fusion dynamics typical of chimpanzees, with no intermediate organisation between dyadic and community relationships, Waibira shows a different organisation with one central subgroup characterized by strong associations (Waibira_A) and a similar number of satellite males with weak associations. Unlike the multilevel organisation observed in other mammal species (e.g., in baboons: Schreier and Swedell, 2009; elephants: de Silva and Wittemyer, 2012, and sperm whales: Whitehead, 2012), not all groupings identified in Waibira contained stable membership or similar patterns of association. Instead, while a single central group of males forms the relatively stable Waibira_A subgroup, with strong inter-individual connections, similar in strength to those found in Sonso at the community level, Waibira_B is a looser aggregation of males whose inter-individual connections are weaker, and who do not spend more time with each other than they do with males from the Waibira_A subgroup. The low modularity in Waibira suggests that the identified subgroups do not represent two distinct and stable cliques but instead highlights that two distinct social strategies may be used by males in this community: a cohesive strategy employed by males in Waibira_A and a satellite strategy, where males form weaker associations, employed by Waibira_B males.

In the first two years of the study period (2015-2017) the Waibira_A subgroup exhibited very stable membership, and six out of nine of these individuals continued to form the core of this subgroup across the four years, in line with the stability observed across two-years in the large Ngogo community’s subgroups (Mitani and Amsler, 2003). However, in the last two years the Waibira_A subgroup showed more variation: appearing to split in the third year and coalesce into a larger subgroup in the last year, drawing in individuals previously associated with Waibira_B. These results suggest substantial nuance in chimpanzee social strategies, such that while some individuals exhibit the same strategy across some or many years, or with major demographic and social shifts, individuals social strategies and therefore the observed social organisation of the community can change. It is unclear what drove individual membership of Waibira_A. Individuals in Waibira_A and Waibira_B were of similar ages and both groups contain a similar range of ranks (although Waibira_A did include the top two ranked males). As we only included individuals who ranged independently across the full four years, we are unable to describe the influence of the next generation of younger males – who became independent within the study period and who were starting to integrate into the network. The larger difference in the composition of the Waibira_A males over the last two-years could therefore be due to an influx of younger males.

Another feature that may impact year-to-year variation in Waibira_A membership is change in the social hierarchy. A change in alpha-ship in the Waibira chimpanzee community occurred shortly before the study period. During the study the current alpha was in Waibira_A and the recently deposed alpha in Waibira_B. Some of the transitions between Waibira_A and Waibira_B may reflect shifts in patterns of association as individuals increasingly shifted their association preferences to the current alpha (Kanngiesser et al. 2011). Our findings highlight the importance of studying higher-order social organisation across the timescales in which social and demographic changes take place for the study species and the integration of subadults into a social network.

Like other large East African chimpanzee communities (Mitani and Amsler 2003) Waibira chimpanzees are able to maintain a larger territory than neighbouring groups - here around double the size of the Sonso community. Subgrouping in other chimpanzee communities appeared to foreshadow a permanent fission between subgroups resulting in two separate communities, either many years (Amsler and Mitani, 2003) or two years (Feldblum et al. 2018) before the split. While in Ngogo individuals from each subgroup overlapped substantially in range (Amsler and Mitani, 2003), in Gombe ranging of the two subgroups quickly diverged over the two-year study period until they exhibited complete separation after fissioning. The two social subgroups in Waibira ranged over highly similar areas within the community territory, suggesting that they are not (yet) in the processes of permanent fission. While increased group size has been correlated to larger territories in several species (van Schaik and van Hooff 1983; Dunbar 1988), including chimpanzees (Lemoine et al, 2020), this is not always the case (Williams et al. 2004), and the availability and distribution of food is also an important correlate of group and territory size (Johnson et al. 2001) and remains to be explored in Waibira and Sonso. Our results support previous observations of the impact group demographics can have on social organisation (Shizuka and Johnson 2020) and provide further support for the suggestion that large groups of male chimpanzees are able to flexibly modify their social organisation within fission-fusion dynamics, while maintaining a single cohesive territory and community (Mitani and Amsler, 2003). Our results suggest that, unlike findings from other chimpanzee communities, any changes in organisation are not restricted to multilevel social organisation or the formation of stable subgroups or cliques.

The differences in social organisation between Sonso and Waibira highlight that chimpanzees, like other social animals, flexibly adapt their social organisation. Doing so offers substantial individual flexibility to respond to short- and long-term changes in local socio-ecology in a way that maximises fitness. Despite the differences in community size and total number of adult males, the average number of males found in a party in Sonso or Waibira at any one time was very similar, supporting the argument that atomistic fission-fusion dynamics allow individuals to manage a day-to-day trade-off between safety in numbers and feeding competition (Aureli et al. 2008; Grueter et al. 2012). Similarly, the total number of males within a cohesive (sub)group appeared to be similar: 11 in Sonso’s single cluster, and nine in Waibira_A’s central cluster. Waibira_A and Sonso community shared the most similar distribution of above chance associations (D dissimilarity), suggesting that around ten males can maintain a relatively stable cohesive unit through atomistic fission-fusion. However, atomistic fission-fusion dynamics appear to be insufficient to manage male-male competition when the number of males in the community is more than double this apparent threshold. The integration of alternative strategies adopted by some individuals to mitigate competition into atomistic fission-fusion societies, may allow larger groups of males to function as a single ‘super-community’ (cf Mitani and Amsler, 2003).

Across taxa, individuals within species employ distinct strategies to avoid competition. One common strategy employed by males with regard to mating competition is satellite or peripheral behaviour (Berec and Bajgar, 2011; Tsuji et al. 1994; Jack and Pavelka, 1997). Satellite individuals may be younger (e.g., Setchell, 2003), lower-ranking (Setchell, 2003; Koprowski, 1993), or smaller (Tsuji et al. 1994) as compared to central males, and often occupy locations at the outskirts of the group home range (e.g. Tsuji et al. 1994) and engage in different mating strategies (e.g. Berec and Bajgar, 2011). Our results provide partial evidence that males in the Waibira_B subgroup may be exhibiting satellite behaviour: they form typically weaker associations with other males, potentially avoiding competitive situations arising from more regular association. However, this strategy did not appear to be fully driven by age or social rank – with a large overlap in both, outside of the top two ranking males. Some female chimpanzees have been suggested to occupy peripheral positions within the community as a strategy to avoid feeding competition (Williams et al. 2002). The large number of independent males in the Waibira community may have led some males (Waibira_B) to employ a similar strategy, using satellite or peripheral behaviour to avoid high intensity competition with the current highest ranked males, while retaining access to feeding patches.

Competition for sexual opportunities in Waibira is particularly high, impacted both by the large number of independent males, but also by a relatively low male:female adult sex ratio (mean sex ratio = 1:1.08) over the study period. Chimpanzee communities are typically female biased (Mitani et al. 1996; Lehmann and Boesch 2008), as is seen in the Sonso community (mean sex ratio M:F = 1:2). As female chimpanzees are not sexually receptive year-round (Mitani, Gros-Louis and Richards, 1996) and experience long interbirth intervals (∼4-5 years; Emery Thompson et. al., 2007), even a female biased adult sex ratio often results in a highly male-skewed operational sex ratio. Thus, the evenly matched adult sex ratio in Waibira likely leads to particularly strong male mating competition for reproductive opportunities. In another chimpanzee community with a male biased sex ratio and similar number of males to Sonso community (N = 12), Fongoli in Senegal, high rates of intracommunity killing were suggested to be the result of increased competition between males (Pruetz et al. 2017). In Gombe, the heavily male biased sex ratio was considered as a major driver of the community split (Feldblum et al. 2018). Intracommunity killings of adults have not been observed (or inferred) in Waibira (Hobaiter, pers. comm.), but flexible social behaviour in the Waibira social structure may offer an alternative strategy to mitigate this competition that is only possible for larger groups. While a direct test of this hypothesis requires longitudinal data, the Waibira social networks are suggestive of this pattern. Rather than forming two subgroups with similarly strong inter-individual relationships, the central subgroup is a cohesive clique, including the current alpha and his closest allies, while the other males are in peripheral orbits, and maintain weaker relationships with others overall.

While some highly peripheral Waibira females are observed less often, and so their sons maybe less well habituated while dependent (increased exposure leads to improved habituation; Bertolani and Boesch 2008; Samuni et al. 2014), all independent males are habituated in both the Sonso and Waibira groups, thus, variation in habituation is very unlikely to explain the variation in social strategy. Kinship plays a role in male-male social association, with maternal siblings more likely to associate and cooperate (Langergraber et al. 2007; Sandel et al. 2020); however, as paternity data for Waibira are incomplete, we are unable to test, as yet, the extent to which association is influenced by relatedness between males.

Male-male relationships are often the focus of descriptions of chimpanzee social organization, but by including only male-male relationships we excluded two thirds of potential dyadic relationships in the community (female-male and female-female dyads). Female behaviour and association with males may also impact male reproductive fitness and behaviour (Langergraber et al. 2013; Reddy and Mitani 2020) and excluding females and younger individuals from analysis may alter male network structures (Fedurek and Lehmann 2017). At a smaller scale, by excluding individuals who were not present throughout the entire study period we may have removed some important relationships that contributed to particular individuals’ position in the social network, and we were unable to see the integration of newly independent males into the subgroups in the later years. Including data from across many years is necessary to explore flexibility in fission-fusion dynamics at a species-relevant scale; however, it also removes nuanced changes to relationships over time - coercing dynamic patterns of association into simple snapshots. While we ensured that overall dyadic association remained stable over the four years, Mantel tests cannot account for small variation in individuals’ dyadic association strength or preferred social partners and our understanding of chimpanzee relationships would likely benefit from exploring more nuanced changes in the structure of communities over time.

Our results are in line with previous research suggesting that flexibility in social organisation is widespread across the animal kingdom (e.g., Schreier and Swedell 2009; Morrison et al. 2019; Stead and Teichroeb 2019; Grueter et al. 2020). We find that chimpanzee communities of different sizes employ atomistic fission-fusion social dynamics but show that there may be upper limits to the number of chimpanzee males able to maintain a community through atomistic fission-fusion dynamics alone, and that stable ‘super-sized’ communities can be formed through the incorporation of flexible social organization with males potentially adopting different social roles (central and satellite). Our results support the findings from another large community of chimpanzees, who showed flexibility in male social organization, however we did not find evidence for stable cliques or modular structures as seen in other communities over a 2-year period (Mitani and Amsler, 2003; Feldblum et al. 2018). Our results suggest that while large chimpanzee communities employ flexible social organisation, they may not all do so in the same way. It has been argued that human social organisation developed from multi-male multi-female groups into the highly structured modular societies, where individuals fill different social roles, seen today, through an intermediate step of single-male core-units (Chapais 2013; Swedell and Plummer 2019). Here we show that a modular society may emerge from multi-male multi-female groups in a different way, and that in doing so it may allow for substantial individual flexibility in the strategies available to maximise individual fitness in a dynamic socio-ecological landscape. While humans do this on a larger scale, in bigger communities, and across greater geographic distances, the diversity and flexibility present in chimpanzee social organisation may be closer than previously described to that found in modern human societies, with individuals exhibiting both cohesive and peripheral social strategies.

## ACKNOWLEDGEMENTS

We thank the Waibira and Sonso field assistants, and all staff of the Budongo Conservation Field Station. We thank the Uganda National Council for Science and Technology, the President’s Office, the Uganda Wildlife Authority, and the National Forestry Authority for permission to conduct the research and the Royal Zoological Society of Scotland for their support of the field-site.

## DECLARATIONS

### Funding

The study employed long-term data collected by the Budongo Conservation Field Station who are supported by the Royal Zoological Society of Scotland.

### Availability of data and materials

The data used for this study are available from the GIThub repository https://github.com/Wild-Minds/Multi-level-social-structure_BudongoChimpanzees

### Code availability

RStudio code used for all data analysis and visualisation is available from the GIThub repository https://github.com/Wild-Minds/Multi-level-social-structure_BudongoChimpanzees

### Author contributions

All authors contributed to the study conception and design. GB, KB, and CH conceived the study; GB, KB, and CH extracted the data; GB, KB, LS, and CH conducted the analyses; GB, KB, LS, KZ, and CH wrote the paper.

## COMPLIANCE WITH ETHICAL STANDARDS

### Conflicts of interest/ competing interest

The authors have no conflicts of interest or competing interests.

### Ethical statement

All data were collected as part of the Budongo Conservation Field Station’s long-term data collection, which is observational only and follows the International Primatological Society’s Code of Best Practices in Field Primatology, as well as all applicable international, national, and institutional guidelines for the care and use of animals. Ethical approval for the use of these data was also granted by the University of St Andrews Animal Welfare and Ethics Committee.

## Notes

https://github.com/Wild-Minds/Multi-level-social-structure_BudongoChimpanzees

### Competing Interest Statement

The authors have declared no competing interest.

