## Supplemental Materials for "Flexibility in the social structure of male chimpanzees (*Pan troglodytes schweinfurthii*) in the Budongo Forest, Uganda"

**Supplementary Materials**

**Table S1** Results from Mantel tests comparing network structure across four study years for the Sonso and Waibira chimpanzee communities

| **Community** | **Years** | **Z** | **P-value** |
| --- | --- | --- | --- |
| Sonso | 2015-2016 | 7.316 | 0.002 |
| Sonso | 2015-2017 | 6.475 | < 0.001 |
| Sonso | 2015-2018 | 6.953 | < 0.001 |
| Sonso | 2016-2017 | 6.754 | 0.002 |
| Sonso | 2016-2018 | 7.27 | 0.003 |
| Sonso | 2017-2018 | 6.465 | < 0.001 |
| Waibira | 2015-2016 | 10.215 | < 0.001 |
| Waibira | 2015-2017 | 5.842 | < 0.001 |
| Waibira | 2015-2018 | 6.535 | < 0.001 |
| Waibira | 2016-2017 | 7.397 | < 0.001 |
| Waibira | 2016-2018 | 8.792 | < 0.001 |
| Waibira | 2017-2018 | 5.087 | < 0.001 |

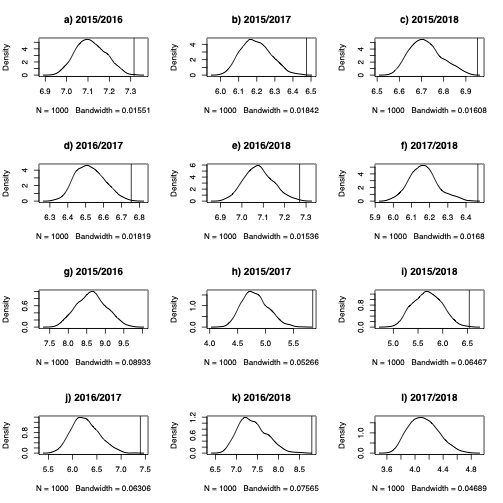

**Fig. S1** Results of Mantel test between four years of the study period. Plots a-f represent yearly comparisons of the Sonso community and plots g-l represent the results from the Waibira community

**Table S2** Network measure results from subset randomisation method used for analyses. P values indicate if observed measure was greater or smaller than expected from null models generated from 10 000 permutations of subsetted dataset.

| Network measure (community) | Observed value | P(greater) | P(smaller) |
| --- | --- | --- | --- |
| Transitivity (Sonso) | 1 | 1 | 1 |
| Transitivity (Waibira) | 1 | 1 | 1 |
| Mean Strength (Sonso) | 3.54 | <0.001 | 0.999 |
| Mean Strength (Waibira) | 3.736 | 0.498 | 0.5021 |
| Modularity (Sonso) | -1.82^-16 | 0.546 | 0.454 |
| Modularity (Waibira) | 0.016 | <0.001 | 0.999 |

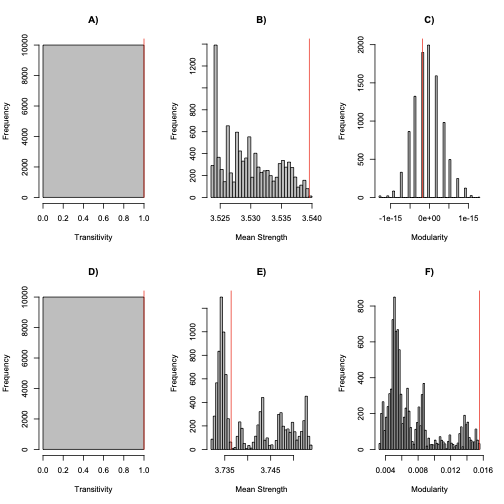

**Figure S2**

Histogram distribution of a) Sonso Transitivity, b) Sonso mean Strength, c) Sonso Modularity, d) Waibira Transitivity, e) Waibira mean Strength and f) Waibira Modularity measures from 10 000 data stream permutations of each social networks using subset randomisation method. Red lines indicate measures from networks created from original party composition data

**Table S3:** D dissimilarity values between communities and between Sonso and Waibira subgroups using subset data. G is the community/subgroup to be compare with G’ (another community/subgroup).

| **G** | **G’** | **D(G, G’)** |
| --- | --- | --- |
| Sonso (Full) | Waibira (Full) | 0.09632188 |
| Sonso (Full) | Waibira_A | 0.05519558 |
| Sonso (Full) | Waibira_B | 0.2014582 |

**Table S4:** results from linear model comparing the individual strength of males in Waibira_A to Waibira_B. In the table the p-value from the model as well as the p-values retained from comparing the observed coefficient to the randomly generated coefficients from 10000 permutations that represent the null model are reported.

| F - statistic (DF[1,20]) | Adjusted R^2 | p-value (model) | P(greater) | P(smaller) |
| --- | --- | --- | --- | --- |
| 20.06 | 0.476 | <0.001 | 0.977 | 0.023 |

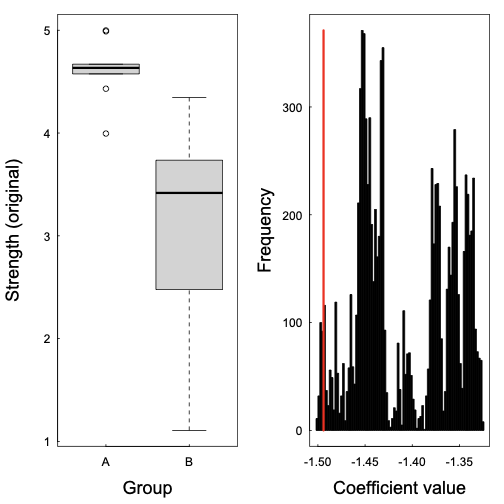

**Figure S3**

Illustration of model results testing the relationship between individual Strength and subgroup membership in the Waibira community subset. a) boxplot illustration of the range of individual Strength in each group with thick lines illustrate the mean Strength in the community, box limits are the lower (25%) and upper (75%) confidence intervals, and the range shows the most extreme (highest/lowest) Strengths that are no more than the range multiplied by the interquartile range. b) the distribution of model coefficients from the null model; the red line indicates the coefficient from the model testing the original data

**Table S5 Levenshtein Distance between Waibira_A subgroups across each pair of consecutive study years in Waibira subset**

*All comparisons are made for the Waibira_A subgroup identified across the four-year study period. Waibira_A was defined as the subgroup with the highest mean Strength. During the 3^rd^ study year two subgroups were identified with similarly high mean Strength so we named these Waibira_A1 (2017/2018a) and Waibira_A2 (2017/2018b). In this table we report the Levenshtein Distance between two Waibira_A subgroups of consecutive years and the number of individuals in the subgroup for each year in parentheses. E.g. between the first two study years the Levenshtein distance between the Waibira_A subgroup was 1 and the subgroup size for both years was 10 individuals, we report this as 1 (10, 10)*

| **Years** | **Levenshtein Distance** |
| --- | --- |
| 2015/2016 – 2016/2017 | 2 (10, 9) |
| 2016/2017 – 2017/2018 | 11 (9, 15) |
| 2017/2018 – 2018/2019 | 10 (15, 13) |

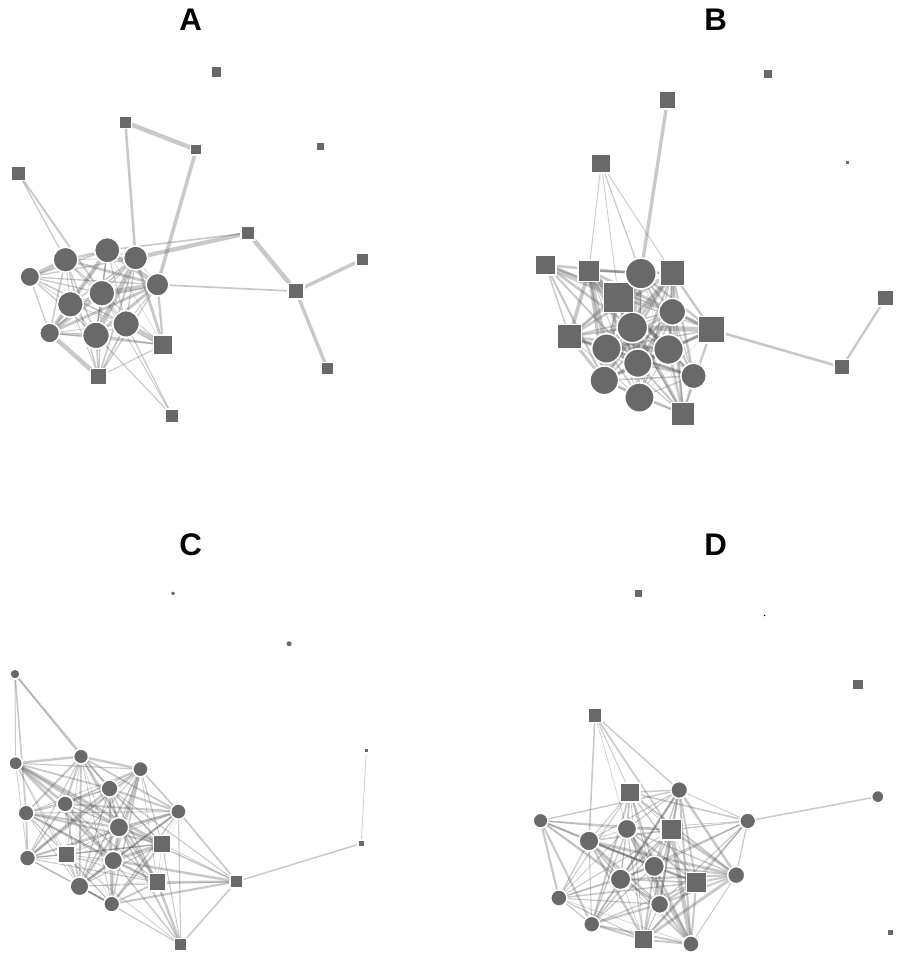

**Figure S4** Sociograms highlighting subgroups identified through Louvain’s clustering algorithm between males in the Waibira community for each of the four years of study: A) 2015-2016, B)2016-2017, C) 2017-2018, D) 2018-2019. Only edges with a value higher than the community mean edges weight of the null models (2015-2016 = 0.17, 2016-2017 = 0.218, 2017-2018 = 0.128, 2018-2019 = 0.156) are present in the network. Node shapes within each network indicate subgroup membership. Node area represents individual Strength multiplied by seven (for clearer visualisation), so that larger nodes have higher individual Strengths. Circular nodes always belong to ‘Waibira_A’ or ‘Waibira_A1’: the subgroup with highest mean Strength

**Table S6** Network measure results from subset randomisation method used for analyses. P values indicate if the observed measure was greater or smaller than expected from null models generated from 10 000 permutations of adult only dataset.

| Network measure (community) | Observed value | P(greater) | P(smaller) |
| --- | --- | --- | --- |
| Transitivity (Sonso) | 1 | 1 | 1 |
| Transitivity (Waibira) | 1 | 1 | 1 |
| Mean Strength (Sonso) | 2.391082 | 0.0056 | 0.9957 |
| Mean Strength (Waibira) | 2.387714 | 0.8531 | 0.1472 |
| Modularity (Sonso) | -2.122597e-16 | 0.7262 | 0.2751 |
| Modularity (Waibira) | 0.01753887 | <0.001 | 1 |

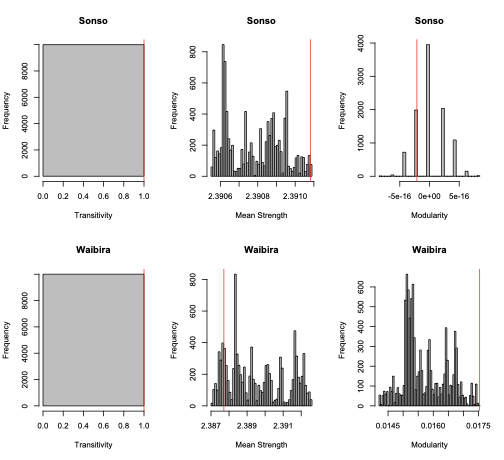

**Figure S5**

Histogram distribution of a) Sonso Transitivity, b) Sonso mean Strength, c) Sonso Modularity, d) Waibira Transitivity, e) Waibira mean Strength and f) Waibira Modularity measures from 10 000 data stream permutations of each social networks using adult data only and individual randomisation method. Red lines indicate measures from networks created from original party composition data

**Table S7:** D dissimilarity values between communities and between Sonso and Waibira subgroups using adult only dataset. G is the community/subgroup to be compare with G’ (another community/subgroup).

| **G** | **G’** | **D(G, G’)** |
| --- | --- | --- |
| Sonso (Full) | Waibira (Full) | 0.3573163 |
| Sonso (Full) | Waibira_A | 0.2304035 |
| Sonso (Full) | Waibira_B | 0.4876596 |

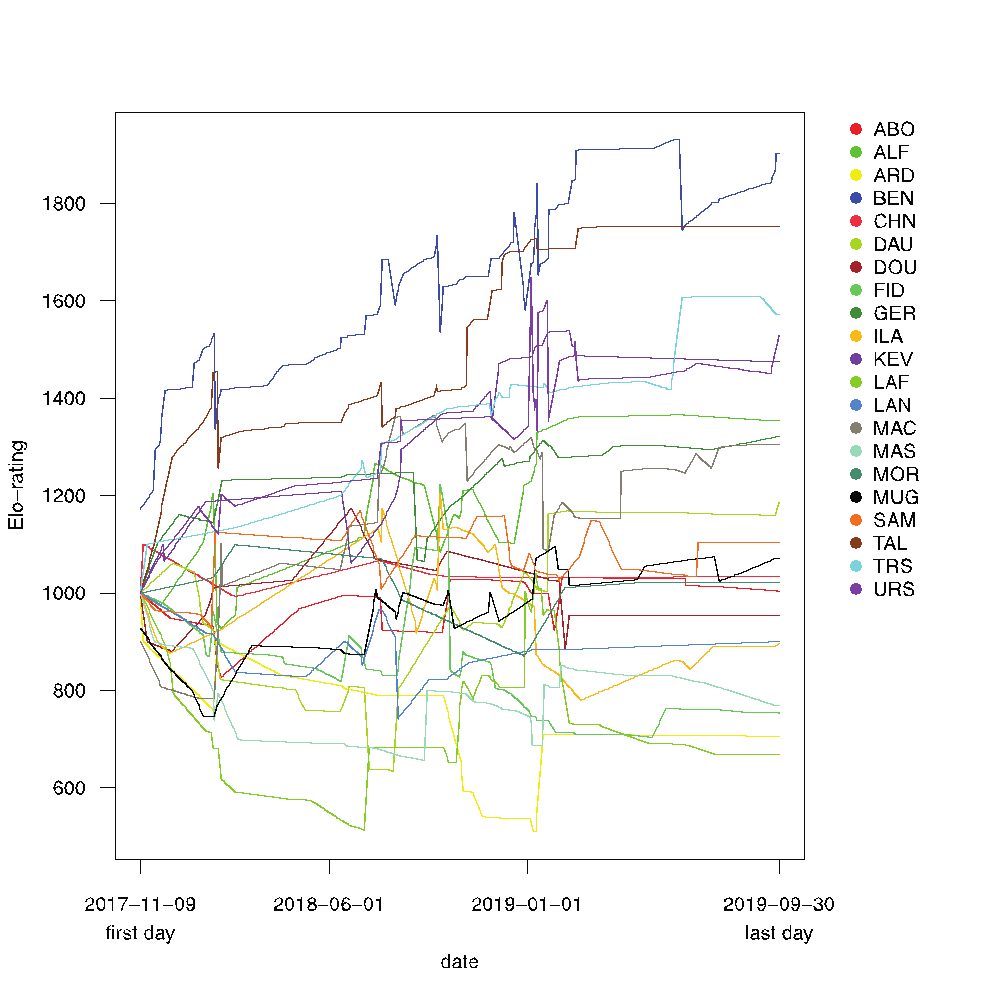

**Fig. S6** Plot showing the rank trajectory of elo-rating over the final two years of the study period for males in Waibira. We could only measure rank over the final two years of the study period because we did not have these data available before then. To calculate ranks we used the package EloRating (Neumann, 2020) in RStudio. We used pant-grunt interactions as a proxy for win-loss interaction where the individual who produced the pant grunt was the loser and the receiver was the winner. We used a starting value of 1000 and k-value of 200 and included in our dataset all pant-grunt interactions that occurred within the community (including those that involved females, and immatures). We then used the final elo-rating score for each individual at the end of the study period to determine their rank (between 1-22) with 1 being the highest ranking and 22 the lowest. Group membership: Waibira A included ALF, BEN, DOU, FID, GER, MAC, MAS, TAL, TRS, and Waibira_B included ABO, ARD, CHN, DAU, ILA, KEV, LAF, LAN, MOR, MUG, SAM, URS. No pant-grunt interactions were observed for one of the Waibira_B members (KAS)

**Table S8** t-test results testing for the difference in home range overlap between dyads within as compared to between subgroups in the Waibira community using only the first party composition scan of each day that was recorded in a known block

| **Proportion of Home Range** | **t** | **p-value** | **Mean within subgroup** | **Mean between subgroup** |
| --- | --- | --- | --- | --- |
| 5% | -0.1 | 0.92 | 0.49 | 0.48 |
| 25% | 1.31 | 0.19 | 0.55 | 0.59 |
| 50% | 1.76 | 0.08 | 0.68 | 0.72 |
| 75% | 1.36 | 0.17 | 0.82 | 0.84 |
| 95% | 1.43 | 0.15 | 0.91 | 0.92 |

**Table S8** t-test results testing for the difference in home range overlap between dyads within as compared to between subgroups in the Waibira community using all party composition scans recorded within known blocks

| **Proportion of Home Range** | **t** | **p-value** | **Mean within subgroup** | **Mean between subgroup** |
| --- | --- | --- | --- | --- |
| 5% | -0.46 | 0.6299 | 0.33 | 0.31 |
| 25% | -1.12 | 0.28 | 0.5 | 0.53 |
| 50% | -1.86 | 0.07 | 0.68 | 0.73 |
| 75% | 1.47 | 0.15 | 0.78 | 0.81 |
| 95% | 1.28 | 0.2 | 0.85 | 0.87 |

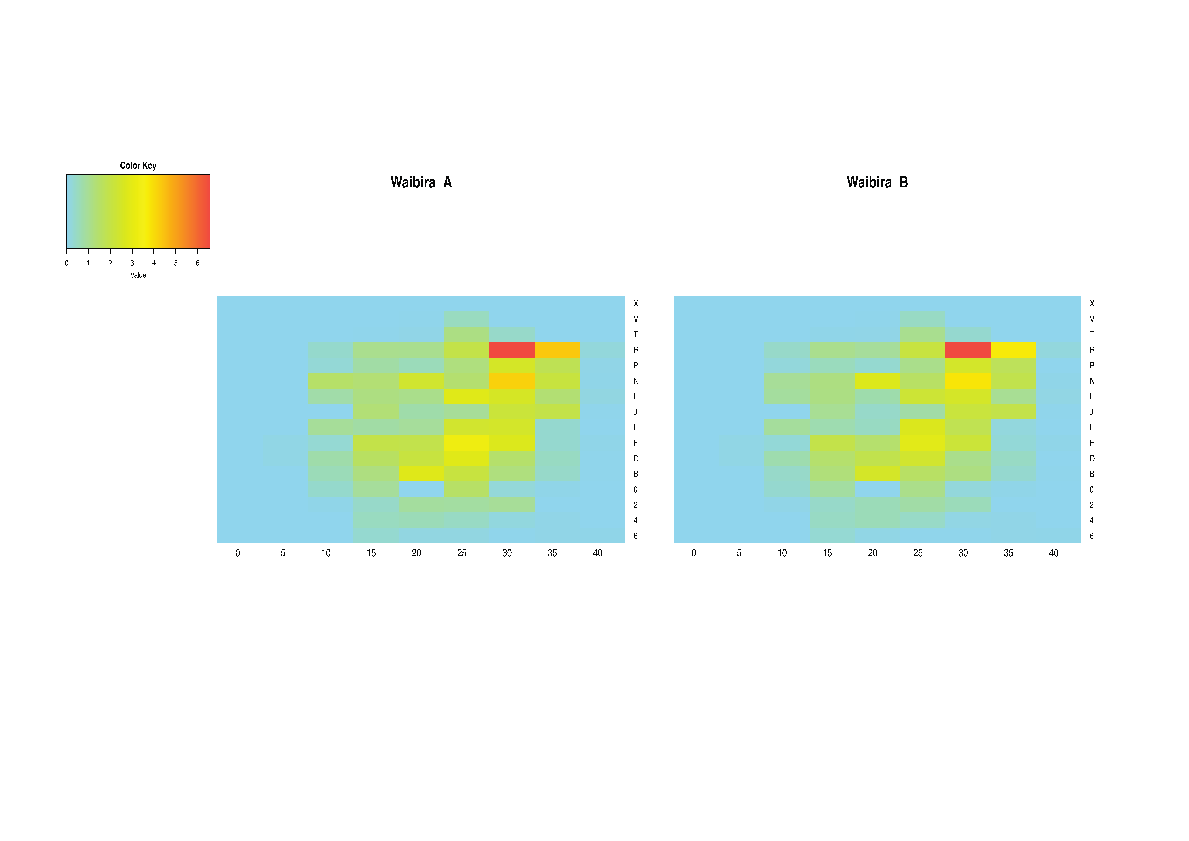

**Fig. S7** Heat maps illustrative the ranging patterns of Waibira_A and Waibira_B from the Waibira community. These maps include all party composition scans and includes all parties in which one or more individuals from the group were present. Colour key indicates the proportion scans were individuals from each group were observed in each block with red indicating the highest proportion time in the block and blue indicating no time spent in the block

**Table S9** Membership and mean Strength of subgroups identified each year. In the membership column each three year code corresponds to one individual ID

| **Year** | **Group** | **Members** | **Mean Strength** |
| --- | --- | --- | --- |
| 2015 | Circle (Waibira_A) | ALF, BEN, DAU, DOU, FID, GER, MAC, MAS, SAM, TRS | 4.79 |
| 2015 | Square | ABO, ARD, CHN, ILA, KAS, LAF, LAN, MOR, TAL | 2.58 |
| 2015 | Triangle | KEV, MUG, URS | 2.61 |
| 2016 | Circle (Waibira_A) | ALF, BEN, DOU, FID, GER, MAC, MAS, SAM, TAL, TRS | 5.62 |
| 2016 | Square | ABO, ARD, CHN, DAU, ILA, KAS, KEV, LAF, MOR, MUG, URS | 3.79 |
| 2017 | Circle (Waibira_A1) | ABO, ARD, BEN, DAU, FID, ILA, LAF, LAN, MUG, SAM, TAL | 2.9 |
| 2017 | Square (Waibira_A2) | ALF, DOU, GER, KAS, MAC, MAS, TRS | 2.72 |
| 2017 | Triangle | CHN, KEV, MOR, URS | 1.48 |
| 2018 | Circle (Waibira_A) | ALF, ARD, DAU, DOU, FID, GER, ILA, LAF, LAN, MAC, MAS, MUG, SAM, TRS | 3.08 |
| 2018 | Square | ABO, BEN, CHN, KAS, KEV, MOR, TAL, URS | 2.42 |
