## Supplementary figures and images for "Flexibility in the social structure of male chimpanzees (*Pan troglodytes schweinfurthii*) in the Budongo Forest, Uganda"

### Supplemental Figure 1

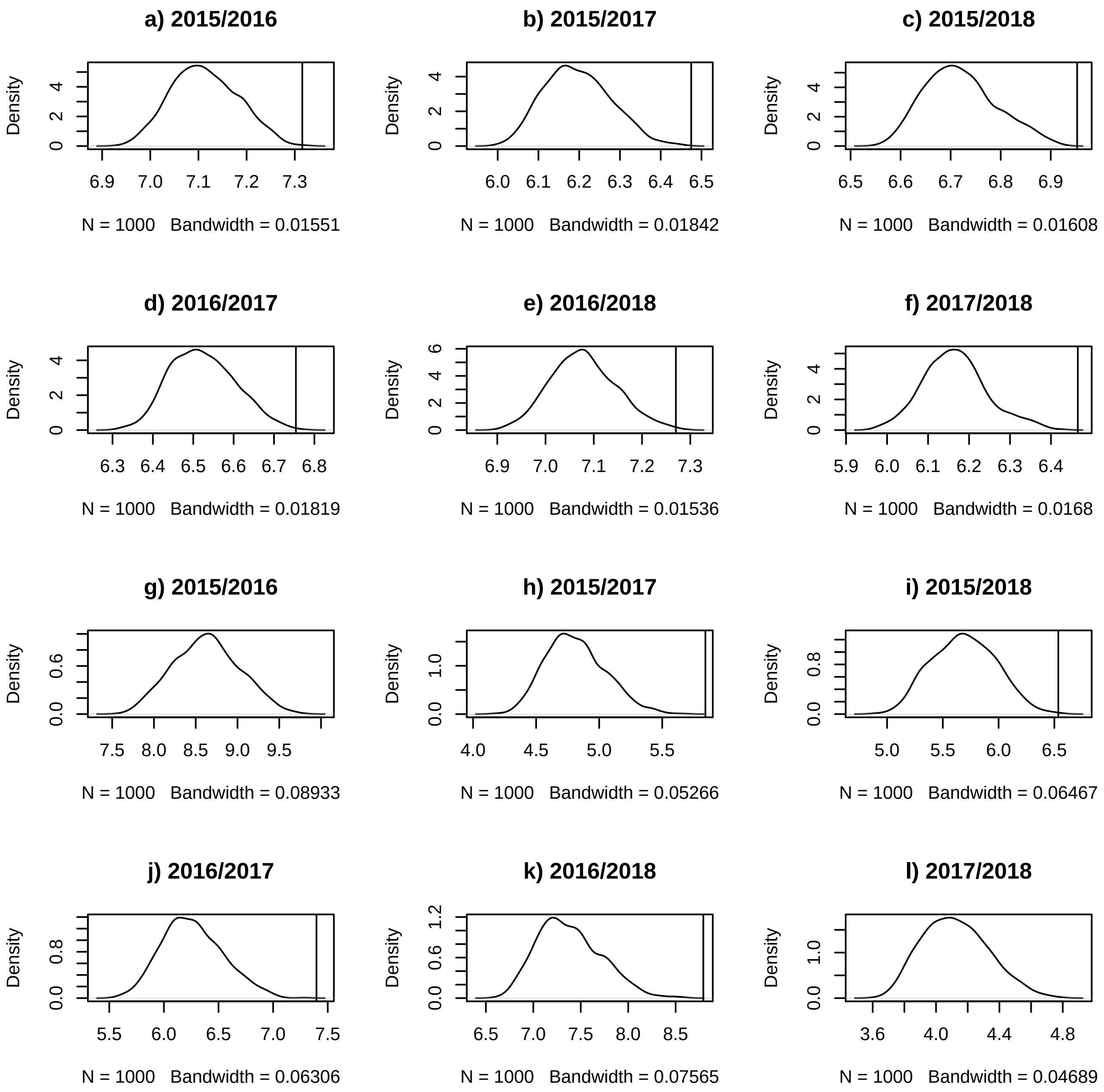

### Supplemental Figure 2

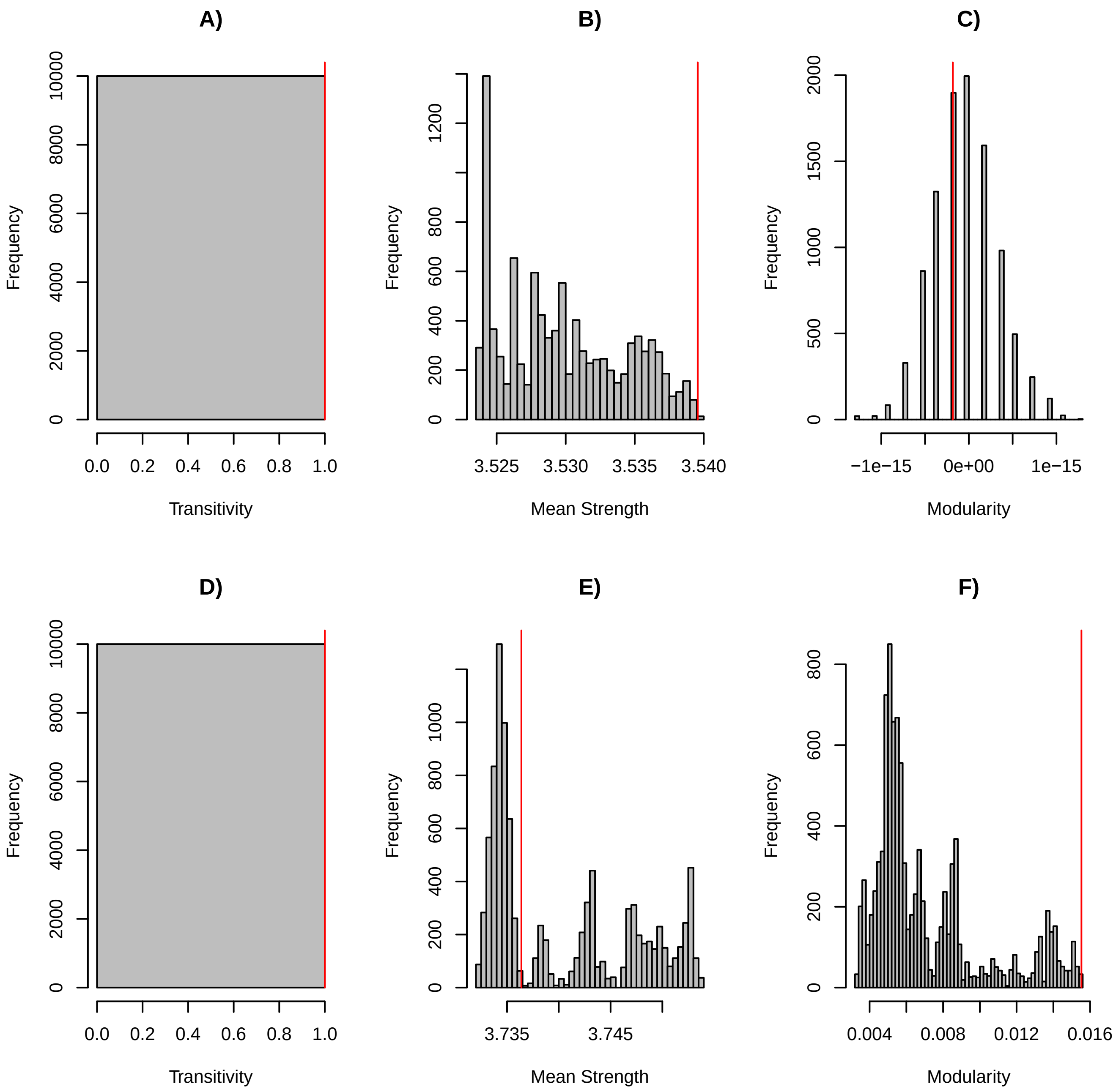

### Supplemental Figure 3

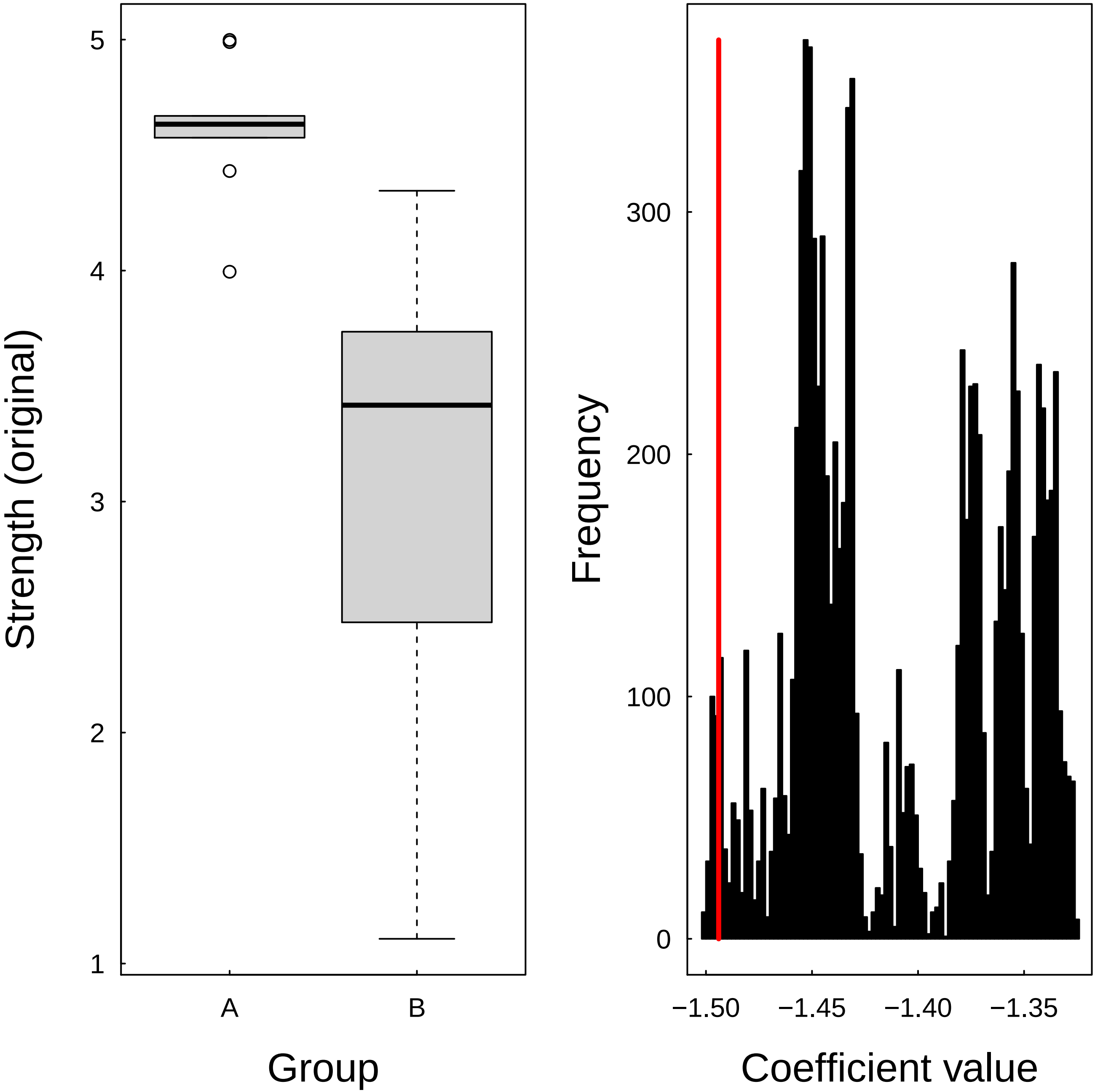

### Supplemental Figure 4

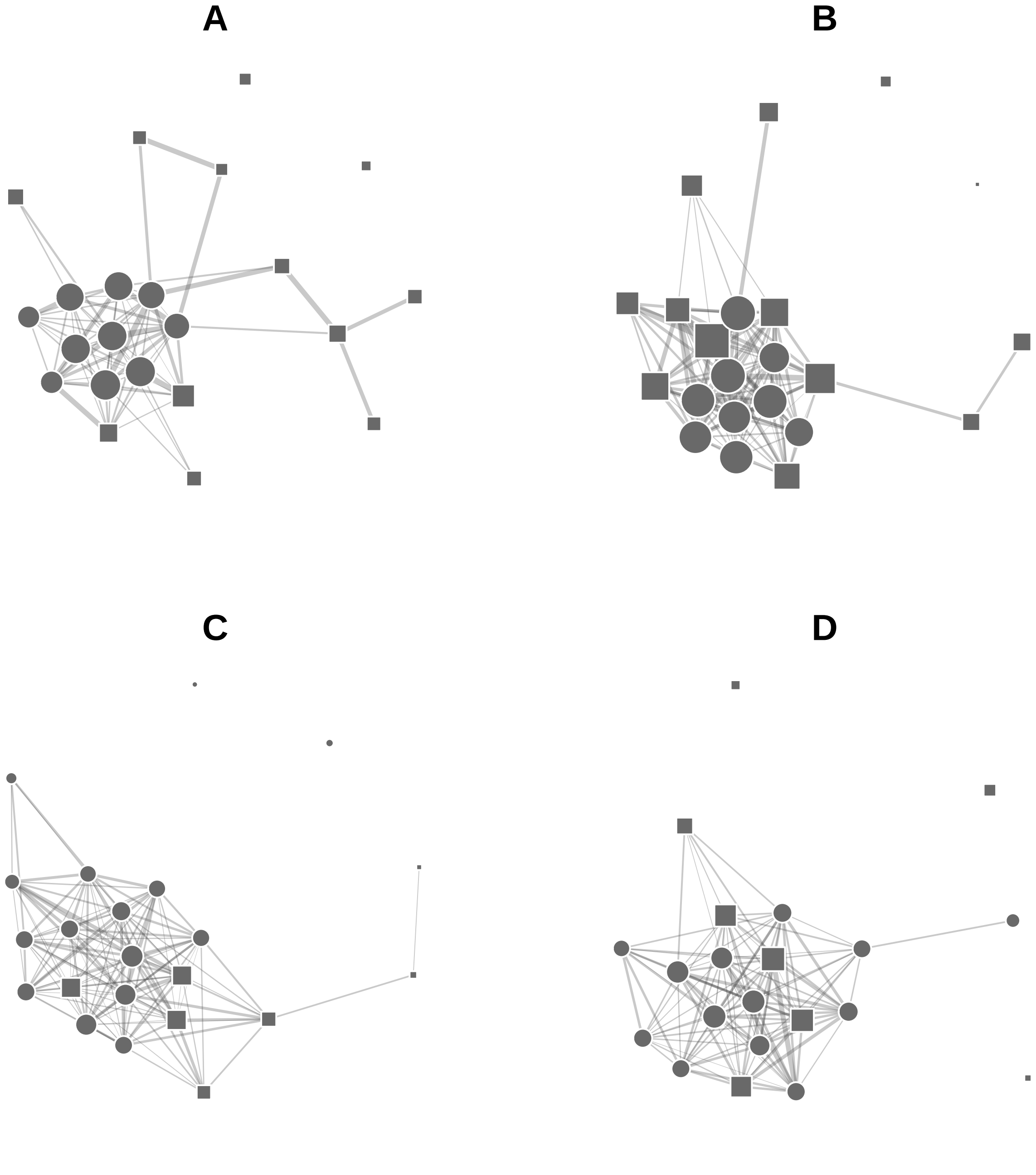

### Supplemental Figure 5

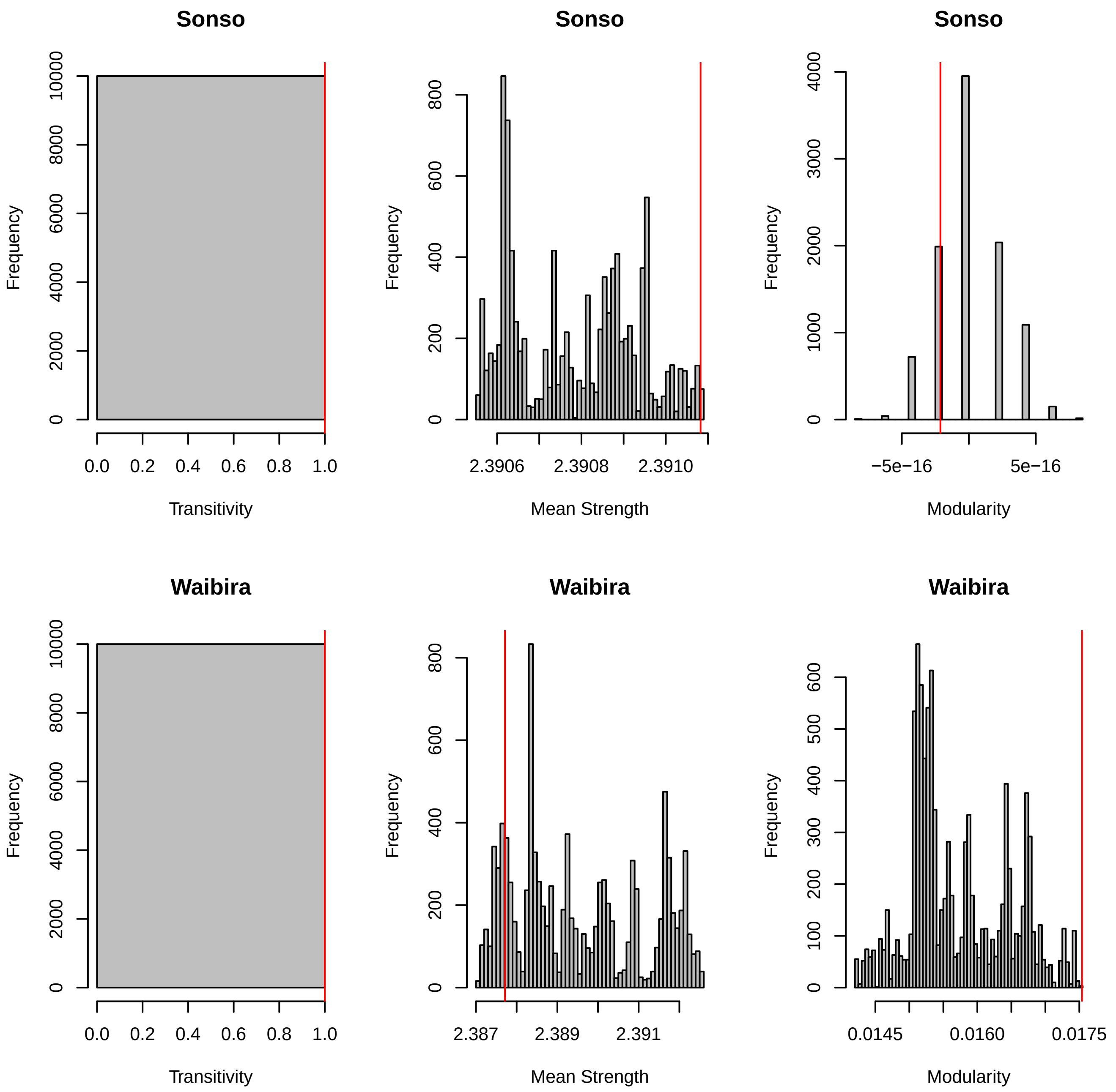

### Supplemental Figure 6

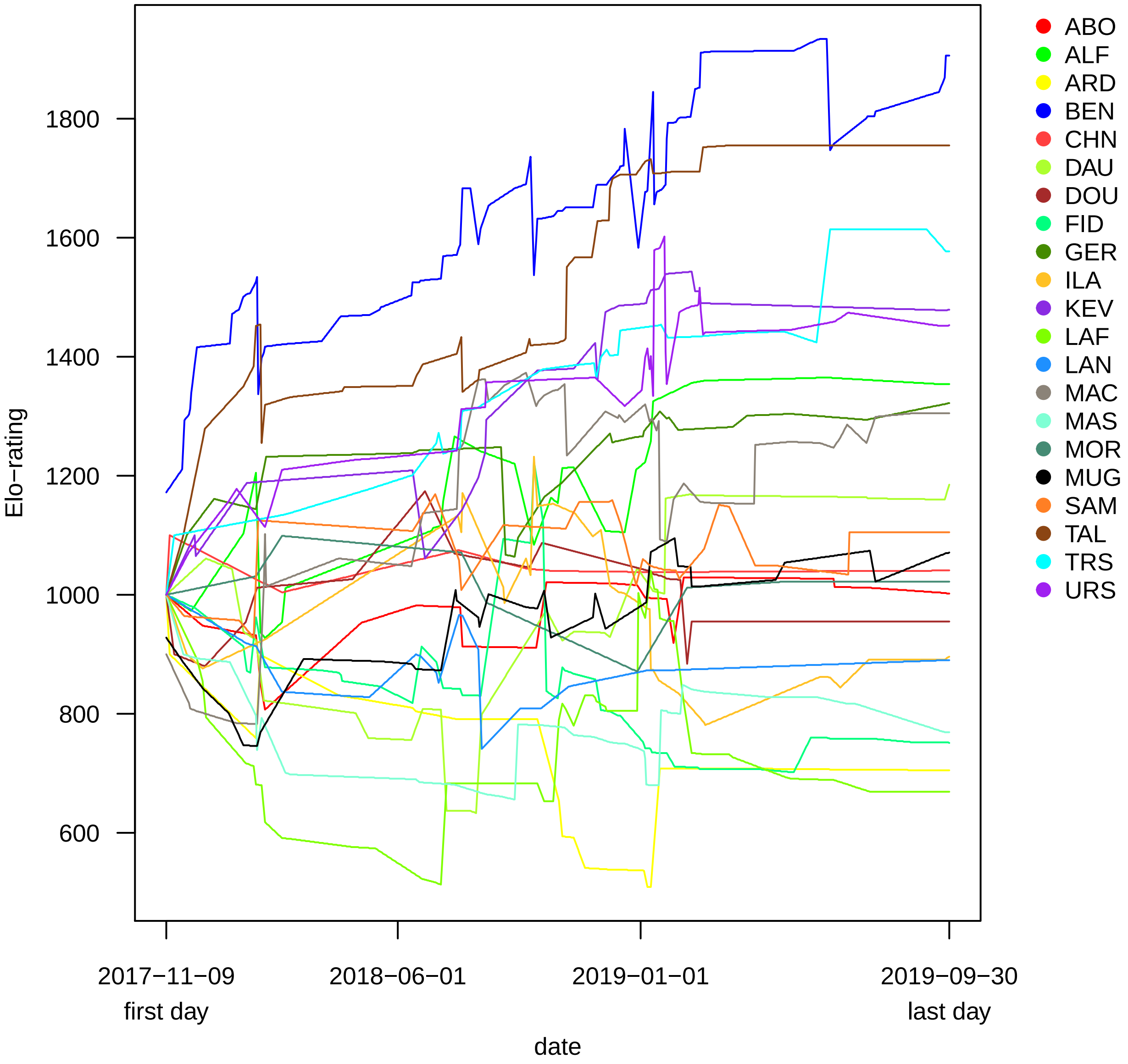

### Supplemental Figure 7

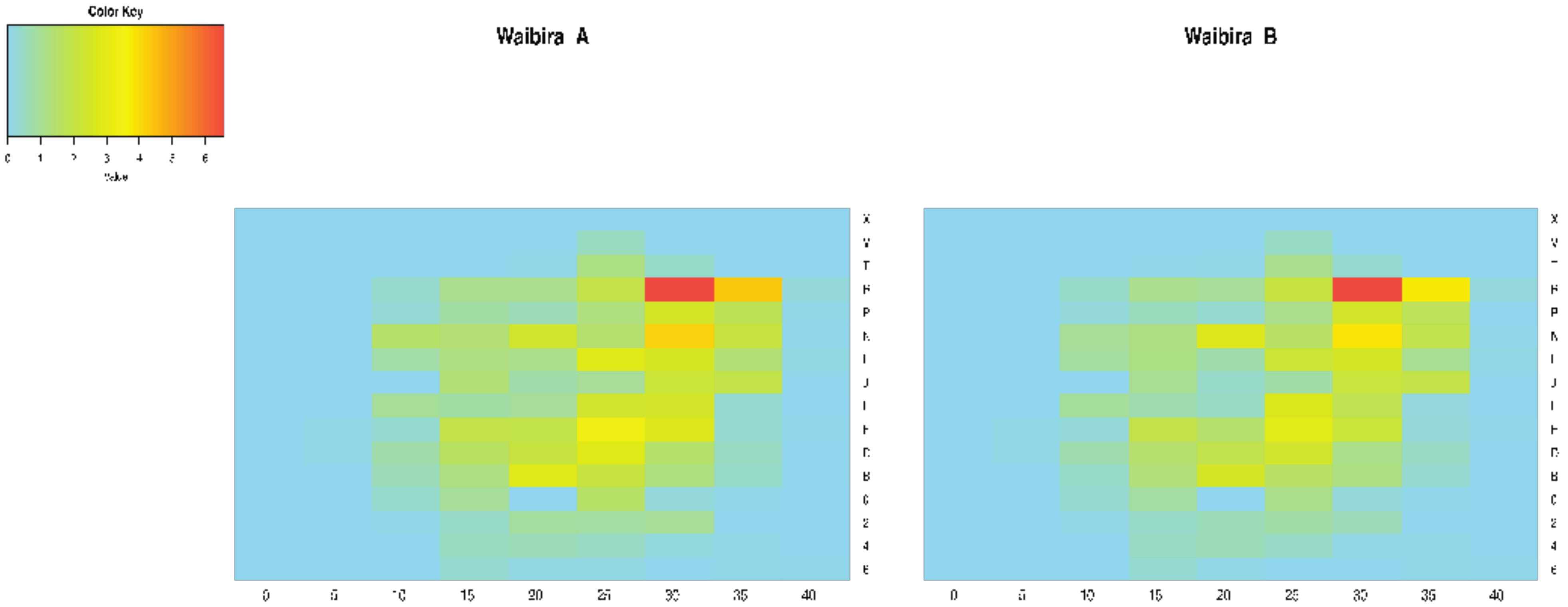
